# Modulation of Polar Auxin Transport Identifies the Molecular Determinants of Source-Sink Carbon Relationships and Sink Strength in Poplar

**DOI:** 10.1101/2023.03.12.532247

**Authors:** Vimal Kumar Balasubramanian, Albert Rivas-Ubach, Tanya Winkler, Hugh Mitchell, James Moran, Amir H. Ahkami

## Abstract

Source-to-sink carbon (C) allocation driven by the sink strength, i.e., the ability of a sink organ to import C, plays a central role in tissue growth and biomass productivity. However, molecular drivers of sink strength have not been thoroughly characterized in trees. Auxin, as a major plant phytohormone, regulates the mobilization of photoassimilates in source tissues and elevates the translocation of carbohydrates toward sink organs, including roots. In this study, we used an ‘auxin-stimulated carbon sink’ approach to understand the molecular processes involved in the long-distance source-sink C allocation in poplar. Poplar cuttings were foliar sprayed with polar auxin transport modulators, including auxin enhancers (AE) (i.e., IBA and IAA) and auxin inhibitor (AI) (i.e., NPA), followed by a comprehensive analysis of leaf, stem, and root tissues using biomass evaluation, phenotyping, C isotope labeling, metabolomics, and transcriptomics approaches. Auxin modulators altered root dry weight and branching pattern, and AE increased photosynthetically fixed C allocation from leaf to root tissues. The transcriptome analysis identified highly expressed genes in root tissue under AE condition including transcripts encoding polygalacturonase and β-amylase that could increase the sink size and activity. Metabolic analyses showed a shift in overall metabolism including an altered relative abundance levels of galactinol, and an opposite trend in citrate levels in root tissue under AE and AI conditions. In conclusion, we postulate a model suggesting that the source-sink C relationships in poplar could be fueled by mobile sugar alcohols, starch metabolism-derived sugars, and TCA-cycle intermediates as key molecular drivers of sink strength.

## Introduction

Source tissues (mainly leaves) fix atmospheric C into carbohydrates and export the photoassimilates to the sink tissues (e.g., roots) (Tegeder and Masclaux-Daubresse 2018; White et al. 2015). Carbohydrate assimilation and partitioning through communication between the source (exporters of photoassimilates) and sink tissues (importers of fixed carbon) have a crucial role in determining plant growth and biomass yield (Lemoine et al. 2013; Bihmidine et al. 2013). Moreover, source-to-sink C partitioning plays a critical role in determining the architecture of a plant, particularly major sink organs and carbohydrate reserves like the root system (Nielsen et al. 1994; Loescher et al. 1990). The source-sink communication that governs this process integrates several internal (source activity and sink demand) and external (water, temperature, and nutrient availability) cues, which requires the co-ordination between different biochemical pathways, genes, and signaling mechanisms.

Sink strength has been defined as the ability of a sink organ to import photoassimilates, which is co-regulated by sink size (the total biomass of sink tissue) and activity (specific uptake rate of the resource) (Ho 1988; Smith et al. 2018). Sink strength is a critical element of dry matter partitioning in the whole plant (Marcelis 1996) and a key to biomass productivity and resilience of yield (Smith et al. 2018; Chamont 1993). Some of the sucrose transport pathways and determinants of carbohydrate storage and utilization in sink organs, including stems, seeds, flowers, and fruits, have been identified in different plant species (Bihmidine et al. 2013). However, the molecular drivers of sink strength have not been thoroughly identified and characterized in root tissue (as a major sink organ) of tree species. Therefore, a comprehensive integrated molecular and physiological analysis is required to understand the relationships between photoassimilates availability and the molecular determinants of sink tissue growth and activity in trees.

Poplar is a prevalent woody feedstock for next-generation biofuels and is acknowledged as the most productive native tree species in the northern hemisphere. Moreover, it is a good target for C sequestration due to its ability to separately control tissue composition above and below ground (Wullschleger et al. 2005). Molecules and mechanisms involved in C allocation in poplar have been investigated (Unda et al. 2017; Coleman et al. 2008; Mayrhofer et al. 2004; Sauter 1988; Witt and Sauter 1994; Zhang et al. 2014). Phloem sap analysis in different tree species, including poplar, identified sucrose, sugar alcohols, and raffinose family oligosaccharides as major abundant metabolites playing important roles in C allocation and sink establishment (Dominguez and Niittylä 2021). In a study by Jeong et al. 2004, metabolites associated with the sink-to-source transition in developing leaves of *Populus tremuloides* (aspen) were identified, among which carbohydrates, amino acids, and other organic acids were the most abundant metabolites. In addition, extensive fluctuations in malate and the malate-to-citrate ratio, as well as a shift in the contents of the primary amino acid asparagine, were observed during rapid leaf expansion, indicating broad metabolic alterations during a short-distance source-to-sink communication in *Populus sp.* (Jeong et al. 2004). However, a comprehensive analysis of the long-distance source (leaf)-to-sink (root) communication process in poplar has not yet been reported.

Phytohormone levels are one of the important factors that regulate source-sink interactions in plants (Li et al. 2020; Roopendra et al. 2018; Roitsch;Ehneß 2000 and Mishra et al. 2022). As a major plant phytohormone, auxin (in a dose-dependent manner) stimulates the mobilization of carbohydrates in source tissues (leaves in the upper shoot) and elevates their translocation toward sink organs, including root (Hansen and Grossmann 2000; Proels and Roitsch 2009; Smith and Samach 2013), thereby regulating the sink strength (Jing et al. 2022). The polar auxin transport (PAT) system comprises several auxin influx and efflux carrier proteins that facilitate the auxin basipetal transport process (Michniewicz et al. 2007; Petrášek and Friml 2009). PAT modulators include synthetic molecules such as polar auxin transport enhancers like Indole-3-butyric acid (IBA) (Damodaran and Strader 2019) and 1-Naphthaleneacetic acid (NAA) (Singh et al. 2009) and polar auxin transport inhibitors like Naphthylphthalamic acid (NPA) (Abas et al. 2021) and 2,3,5-triiodobenzoic acid (TIBA) (Amijima et al. 2014). These compounds have been commonly used as toolkits to study the hormonal influence on plant physiology and development and have been shown in many plants and woody species to enhance or repress the sink tissue growth and activity (Vielba et al. 2020; Ahkami et al. 2013; Guan et al. 2020; Garrido et al. 2002; Lin and Sauter 2019; Smulders et al. 1988; Copes and Mandel 2000; Taylor and Hoover 2018; Lu et al. 2017; Jing et al. 2022). However, these synthetic auxin modulators have not been applied in a foliar manner to monitor long-distance C relatioships between leaf (source) and root (sink) tissues.

In this work, using an “auxin-stimulated carbon sink” approach, we studied the molecular processes involved in the long-distance source-sink C allocation in growing young poplar plants. We hypothesized that enhancement or repression of phloem-transported auxin levels could modulate C allocation and potentially reveal key genes and metabolites that can act as molecular drivers of sink strength in poplar. To this end, we treated poplar cuttings with a foliar spray of AEs (IBA and IAA) and AI (NPA), followed by a comprehensive analysis of poplar leaf, stem, and root tissues using integrated biomass evaluation, root phenotyping, C isotope labeling, metabolomics, and transcriptomics approaches. We identified several upregulated transcripts in root tissue under AE condition that could contribute to increasing sink size and activity. Further, the metabolic profiling of plants treated with auxin modulators revealed altered relative abundance levels of specific metabolites like sugars, sugar alcohols, and TCA cycle intermediates in roots which could satisfy the C demand during root growth in poplar. Altogether, this work highlights the molecular aspects of the source-sink regulatory mechanisms, which are scarcely known in trees, including the molecular drivers of sink strength.

## Materials and Methods

### Plant growth conditions and treatments

*Populus tremula* × *Populus alba*, clone INRA 717-1B4 one-year-old woody stem cuttings with 6-8 dormant buds were provided by Dr. Stephen DiFazio (West Virginia State University) and stored at −80℃ until use. One week before the start of planting, the woody cuttings were transferred to 4℃. Followed by that, the woody cuttings were planted in Pro-mix BX soil (Promix, PA, USA), and kept in a walk-in growth chamber with a temperature of 24 °C /18 °C (day/night), a light intensity of 400µM/sec and a photoperiod of 16h/18h (day/night). The poplar cuttings were fertilized once per week using 100 ppm of Jack’s professional fertilizer (Jr Peters Inc, PA, USA). After approximately one month, they were transplanted to bigger soil pots (4”x4”x9.5”) to facilitate root growth and grown for an additional one month. Once shoots were formed on the main woody stems, leafy cuttings with 4-5 inches in size, were excised from the apical shoot tips and the internodes below apical tips. Each cutting had just one leaf left on them to reduce transpiration. The stem base of all cuttings was treated with commercially available rooting powder (Rhizopon AA#2, Hortus USA Corp., NY, USA) and planted in the Pro-mix BX soil in 4” square pots to enable the rooting process. The cuttings were grown under similar growth conditions as described above. After 21 days, the leafy cuttings were checked for newly formed roots and then transplanted onto 4”x4”x9.5” pots containing Pro-mix BX soil and fertilized. The cuttings were allowed to grow for an additional two weeks and the newly developed leaves in the 4-5 weeks old cuttings were used for auxin foliar spraying experiments. For Auxin enhancers (AEs), 3 µM NAA (α-Naphthalene Acetic Acid) and 4.5 µM IBA (Indole-3-Butyric Acid) were mixed and sprayed as an enhancer cocktail while for auxin inhibitors (AI), 100 µM NPA (Naphthylphthalamic Acid) was used. AE and AI were prepared in 1N KOH (control solution) as per manufacturer recommendation (Phytotechnology Laboratories Inc, KA, USA). Since there were no earlier reports of foliar spraying of these auxin modulators in poplar, we optimized the spraying concentration based on (i) our previous experiences in foliar spraying in other plant species like *Petunia hybrida* (Ahkami et al. 2013) which served as our base concentration, and (ii) published literatures on woody plants where NPA (auxin inhibitor) were directly sprayed at the root initiation site which served as our upper limit concentration (Valladares et al. 2020). We were mindful of the fact that very high concentrations of these synthetic auxin modulators can result in chemical stresses on source leaves with potential reduction in plant biomass development, as reported in other studies (Nassef and El-aref 2018; Di Benedetto et al. 2015; Rastogi et al. 2013). Based on these factors, in our pilot experiment, we optimized AE and AI concentrations that resulted in changes in root phenotype but not altered source leaf growth in terms of the number of leaves, leaf greenness, and dry weight. The control plants were sprayed with 1N KOH solution which grew normally and established leaf growth regularly at the end of the spraying experiment (supplementary table S1). Therefore,1N KOH caused no negative phenotypic effects on leaves. The leaves of a minimum 6-8 rooted cuttings were sprayed between 10:00 AM and 11:00 AM, with AE and AI agents for a period of 11 days. We adopted 11 days as a treatment timeline for this study based on a pilot experiment, in which we determined that this duration resulted in a measurable difference in root phenotypes between AE and AI, without any detrimental effects on foliar tissues. For the foliar spraying, we used 8, 6, and 8 rooted cuttings for control, AE, and AI treatments, respectively. Each treated plant was sprayed with 5 mL of AE or AI reagents for the first 4 days, followed by 6.2 mL for the consecutive 2 days, and 7.4 mL until plant harvest. After 11 days of foliar spraying, plants were carefully uplifted, soil was removed by gentle washing in water, and leaf, stem, and root tissues were harvested and flash-frozen in liquid nitrogen. The various tissue samples were kept at −80 °C until being processed for omics analyses. A separate batch of hormone-sprayed plants was harvested and dried at 60℃ for two days and used to measure dry biomass weights. All sample collections for downstream molecular works were performed between 10:00-11:00 AM (4-5 hours post dawn).

### Root imaging and data analysis

The hormone foliar-sprayed plants along with controls were used for root imaging, in which the roots were washed to remove soil and placed in a water tray with black cloth submerged beneath the roots to create a contrasting background. For root image analysis, we had 6, 7, and 8 rooted cuttings for control, AE, and AI treatments, respectively. The roots were manually separated using forceps, so that all root branches were well displayed. Images were captured using a digital camera placed on a tripod. The root pictures were then analyzed using the DIRT platform (http://dirt.cyverse.org/)(Bucksch et al. 2014, Das et al. 2015). A circular scale marker was placed in each picture to adjust for the camera zoom variations between different pictures. Briefly, the root images were uploaded into the database, and the thresholding of roots was performed by selecting the appropriate parameter. Then, scale values were entered, root system architecture (RSA) parameters were selected, image analysis was performed, and an output file with data was exported. Selective parameters from the output file, such as lateral root branching frequency (no. of lateral roots/nodal root path length), root area (total area of the roots), and maximum root width (maximum of the calculated width in the width height diagram) are presented in Fig. 2.

### ^13^CO_2_ labeling experiment

For the isotope labeling experiment, due to the technical difficulty associated with performing a simultaneous isotope enrichment alongside foliar hormone spraying, we first applied the foliar spray to plants with AE and AI modulators, and then transferred the treated plants to a chamber for isotope enrichment (Fig. 1). We had 4 biological replicates for controls, AE and AI treatments. The foliar hormone sprayed plants and control plants were moved into a transparent Plexiglass chamber for stable isotope labeling using ^13^CO_2_ (99+ atom%, Sigma Aldrich, St. Louis, Missouri, USA). This chamber was incubated within a Conviron walk-in plant growth chamber (model no. GR48, Winnipeg, Manitoba, Canada), which was maintained at 24 °C during the day and 18 °C at night using a 16 h: 8 h day/night cycle. We injected ^13^CO_2_ through a septum port built into the labeling chamber, daily over a four-day incubation period. The plants were exposed to a ^13^CO_2_ enriched headspace for four days, where we sought to maintain total CO_2_ concentrations below ∼800ppm to avoid any artifacts associated with elevated CO_2_ levels.

**Figure 1.**
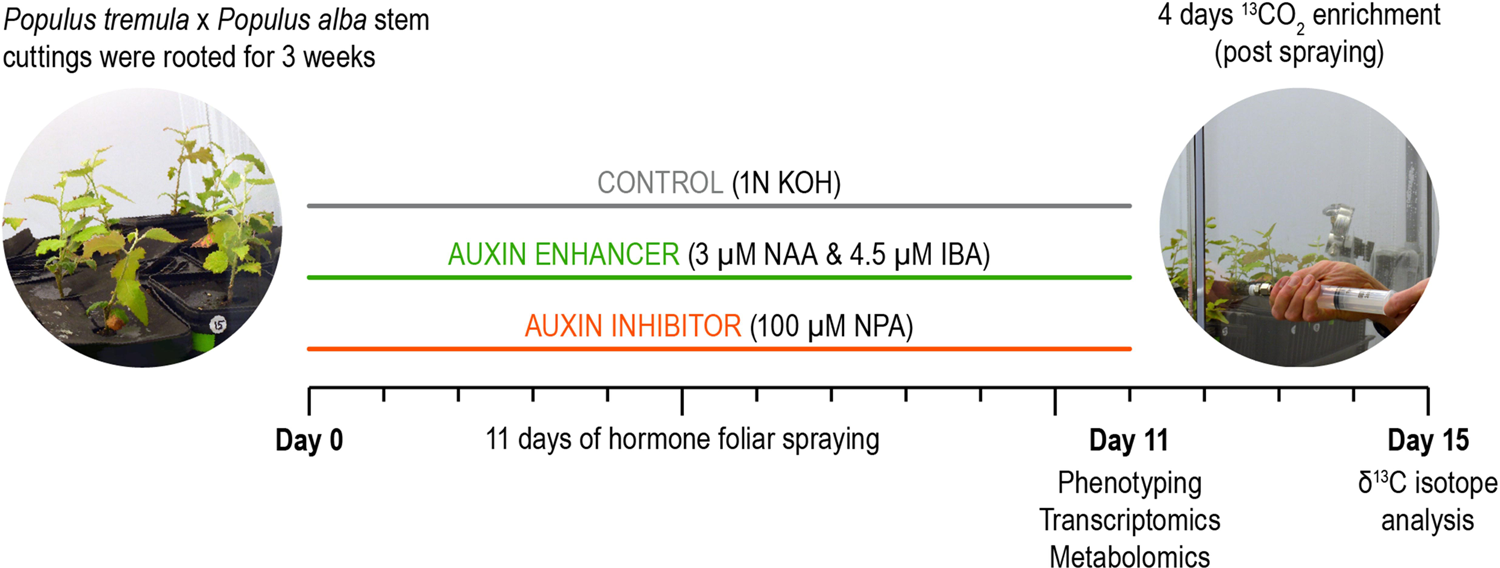
Experimental design and workflow for foliar spraying of auxin modulators, ^13^CO_2_ enrichment, and sample harvesting for omics analyses.

Following ^13^CO_2_ labeling, we removed the plants from the labeling chamber and harvested leaf, stem, and root tissues. Samples were lyophilized, ground to a fine powder using a mortar and pestle, and subsequently aliquoted into tin capsules for isotope analysis. We used a Costech Analytical Elemental Analyzer (Valencia, CA, USA) coupled to a Thermo Scientific Delta V plus isotope ratio mass spectrometer (EA-IRMS; Bremen, Germany) for measuring ^13^C content and all values are reported in delta (δ) notation:

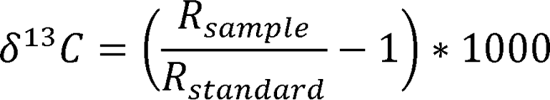

where R_sample_ is the ^13^C to ^12^C ratio of the measured sample and R_standard_ is the ^13^C to ^12^C ratio of Vienne Pee Dee Belmnite. We calibrated the results using a 2-point data correction with in-house glutamic acid stable isotope standards having δ^13^C of −11.09‰ and 16.73‰, where these values were previously calibrated against USGS 40 and USGS 41 (δ^13^C of −26.39‰ and +37.63‰, respectively). All δ^13^C analyses were performed using analytical triplicates.

### LCMS and GCMS-based metabolite profiling

Lyophilized samples were ground using a Qiagen tissue lyzer, and powdered samples were kept at −80ᵒC until ready for metabolite extraction. Polar and semi-polar metabolites were extracted following t’Kind et al. (2008) with minor modifications. Briefly, for each sample, 40 mg of dried powder was added into a clean 2mL glass vial, and 1ml of ultrapure methanol: water (80:20) was subsequently added. Samples were shacked at 1,200 rpm in a Thermomixer (Thermo Fisher Scientific, Waltham, Massachusetts, USA) for 1 hour at 21 °C. Samples were then centrifuged at 13,000 × g for 10 min, and clean supernatants were collected and kept in 2mL glass vials at −80 °C until mass spectrometry analyses.

The total biological replicates used for metabolomic profiling were as follows:

– Leaf LCMS: AE (n=6) and control (n=5); AI (n=5) and control (n=4)
– Leaf GCMS: AE (n=6) and control (n=4); AI (n=4) and control (n=4)
– Stem LCMS: AE (n=4) and control (n=4); AI (n=4) and control (n=3)
– Stem GCMS: AE (n=5) and control (n=4); AI (n=3) and control (n=4)
– Root LCMS: AE (n=5) and control (n=4); AI (n=4) and control (n=3)
– Root GCMS: AE (n=4) and control (n=3); AI (n=3) and control (n=3)

### GC-MS analyses

Gas chromatography-mass spectrometry (GC-MS) analyses were performed on derivatized extracts, for that, 0.5 mL of clean extracts were transferred into clean 2mL glass vials and completely dried in a centrifugal vacuum evaporator. Derivatization of functional groups was performed in two separate steps: methoxymation and silylation. Methoxymation of metabolites was performed using methoxyamine in pyridine solution (30 mg/mL). Each dried sample received 20 ul of methoxyamine solution, and samples were subsequently incubated in a Thermomixer at 1,200 rpm (Thermo Fisher Scientific, Waltham, Massachusetts, USA) for 90 min at 37 ᵒC. This step blocks the carbonyl groups, such as aldehydes, ketones, and carboxylic acids, preventing multiple chromatographic peaks of certain compounds, as sugars, and help to prevent alpha-keto acids from decarboxylation. After the first incubation, silylation of metabolites was performed by adding 80 μl of MSTFA (N-Methyl-N-(trimethylsilyl) trifluoroacetamide) to each of the vials. Samples were thus incubated at 1,200 rpm for 30 min at 37ᵒC. This step mainly derivatizes amides, amines, carboxyl, and hydroxyl groups to trimethylsilyl groups [−Si(CH_3_)_3_]. All derivatized extracts were finally vortexed for 10 seconds and centrifuged at 8,000 rpm for 5 minutes. Supernatants were transferred into clean glass vials with 200 µL inserts.

GC-MS of the samples was performed with an Agilent GC 7890A equipped with a HP-5MS column (30 m × 0.25 mm × 0.25 μm; Agilent Technologies) coupled to a MSD 5975C mass spectrometer (Agilent Technologies, Santa Clara, CA). The injection volume was 10 µL in split-less mode and the injection port temperature was set at 250 °C. The chromatographic gradient was set as follows: 1 min constant at 60 °C, the temperature increased constantly to 325 °C at a rate of 10 °C/min (26.5 min ramp) and maintained for 10 min. For the calculation of the retention index (RI; a normalized measure of the retention time), a mixture of fatty acid methyl esters (FAMEs; C8-C28) was injected at the beginning of the sequence. An experimental blank, consisting of methanol: water (80:20) went through all the extraction and derivatization steps, was injected every 15 samples, and was used to determine the instrument background during the data filtering procedures.

### LC-MS analyses

Liquid chromatography-mass spectrometry (LC-MS) analyses were conducted on a Vanquish ultra-high performance liquid chromatographer (UHPLC) (Thermo Fisher Scientific, Waltham, Massachusetts, USA) coupled to LTQ Orbitrap Velos high-resolution mass spectrometer (HRMS) equipped with a heated electrospray ionization source (HESI) (Thermo Fisher Scientific, Waltham, Massachusetts, USA). The metabolites were separated in the LC using a Hypersil gold C18 reversed-phase column (150 × 2.1 mm, 3µ particle size; Thermo Scientific, Waltham, Massachusetts, USA) held at 30 ᵒC. Before analyses, mobile phases were filtered and degassed for 20 min in an ultrasound bath and consisted in water 0.1% formic acid (A) and acetonitrile/water 0.1% formic acid (90:10) (B). The injection volume was set at 5 µL and the flow was maintained constant at 0.3 mL per minute along all the chromatographic gradients. The elution gradient was initiated at 90% A (10% B) and was maintained for 5 min before the gradient changed to 10% A (90% B) until 20 minutes of the chromatography. Those conditions were held for 2 more minutes, and the starting proportions (90% A; 10% B) were recovered over the following 2 minutes. The column was thus washed and stabilized for 11 additional minutes before analyzing the next sample. All samples were analyzed in negative (-) and positive (+) ionization modes. The HRMS (high resolution mass spectrometry) operated at a resolution of 60,000 in Fourier Transform Mass Spectrometry (FTMS) full-scan mode detecting ions between 50-910 m/z. As in GC-MS analyses, blanks were injected every 8 samples to determine the instrument background signal.

### Processing of GC-MS RAW files

GC-MS raw files were processed in Metabolite Detector 2.5 (Hiller et al. 2009). First, raw “.D” files from Agilent were converted to “.CDF” format in Agilent Chemstation. Raw.CDF files were imported in Metabolite Detector and generated “. CDF.bin” files which were used for processing. GC-MS files were deconvoluted, aligned and peaks were matched against an updated version of the library FiehnLib (Kind et al. 2009) with over 850 compounds before the GC-MS dataset was exported to a CSV file. Briefly, RIs for each detected metabolite were calculated using the FAMEs mixture (Fatty Acid Methyl Esters Standard Mixture) and chromatograms were deconvoluted and aligned. Metabolite identification was conducted by matching MS spectra and RI with the FiehnLib library with probability-matching threshold set at 60%. Features matched to metabolites were subsequently validated by matching the measured fragmented MS spectra to NIST14 GC-MS library. Metabolite detector parameters and GC-MS metabolite matching information are detailed in Table S4D.

### Processing of LC-MS RAW files

LC-MS RAW files were processed in MZmine 2.51 (Pluskal et al. 2010). Samples were analyzed for each ionization mode separately. Briefly, chromatograms were baseline corrected, exact masses (MS1) were detected, and ion chromatograms were generated. Ion chromatograms were thus deconvoluted, and individual features were aligned, gap-filled, and matched against our in-house metabolite library according to exact mass and retention time (RT) values. Our in-house metabolite library includes the exact m/z values of proton adducts in both positive (H^+^) and negative (H^-^) ionization modes and the RTs of over 600 standards from primary and secondary plant metabolism. Datasets for each ionization mode were thus exported to a CSV format file and included the aligned metabolomic profiles of all analyzed samples. MZmine parameters details are shown in Table S4E. Metabolite identification based on exact mass and RT corresponds to a second level of putative identification (Sumner et al. 2007). However, the high mass accuracy achieved by HRMS together with highly reproducible RT values substantially reduce the amount of incorrect compound assignations. RT and m/z values of LC-MS metabolite identifications are shown in Table S4E.

### GC-MS and LC-MS dataset filtering

The GC-MS and LC-MS (negative and positive ionization modes) datasets were separately filtered at *cell* level. A *cell* of the dataset is defined by all the replicate samples within the same experimental group, in other words, the group of samples from the combination of all levels between the different categorical factors of the experimental design (tissue and auxin treatment). In metabolomic experiments, AE and AI treatments were performed independently with their own controls for each tissue, separately. Therefore, our metabolite dataset contains 12 different cells: AE-leaf, AI-leaf, control-AE-leaf, control-AI-Leaf, AE-stem, AI-stem, control-AE-stem, control-AI-stem, AE-root, AI-root, control-AE-root, and control-AI-root. The main purpose of filtering the datasets is to significantly reduce instrument background noise and non-representative features (i.e., features detected solely 1 or 2 samples) from the datasets before proceeding with statistical analyses. Those statistical “artifacts” can surge from different steps ranging from the mass spectrometry instrument analyses to the feature detection and aligning steps. Data filtering was performed through 4 primary steps:

1. *Blank threshold.* For each of the detected and aligned metabolomics features (continuous variables), the number of blank samples with positive data values was counted. If data for a given feature was shown in a single experimental blank, it was considered as zero and not considered to be a representative of noise data
2. *Signal to noise (S/N).* S/N was calculated for each *cell* independently. The noise level of each feature was determined from the 6 blank samples. The metabolite features with S/N < 10 across all *cells* were removed from the dataset.
3. *Cell-Zero filter.* To avoid spontaneous data in a given variable, which typically are shown in very low relative abundance, when a *cell* had <30% of replicates with detected data (only 1 replicate), all values for that *cell* were considered zero.
4. *Minimum data*. Metabolite features with data in less than 70% of the replicates across all *cells* were removed from the dataset, and only those variables with at least one *cell* containing data in 70% or more of its replicates were maintained.

### KEGG pathway analysis of significant metabolites identified by untargeted metabolic analysis

The significant metabolites, defined as differentially abundant metabolites under hormone-sprayed condition compared to control condition, identified through our untargeted metabolome analysis were searched in the KEGG pathway database using the KEGG IDs and mapped onto metabolite pathways using the online pathway mapper tool. KEGG maps of individual pathways (e.g., carbohydrate metabolism, secondary metabolism) that contain our query metabolite list were used to reconstruct a comprehensive metabolic map using Affinity designer application for each condition/tissue that were presented in supplementary Figs. S4A-C.

### Transcriptome analysis

The 11-day hormone-treated and control plant tissues were harvested between 10:00 am to 11:00 am (4-5 hours post-dawn). Leaf, stem, and root samples were frozen in liquid nitrogen and ground into a fine powder using mortar and pestle. Total RNA was extracted from 100mgs of tissue powder using GeneJet Plant RNA purification mini kit (Thermo scientific, USA), and the concentration of RNA was measured using a nanodrop (Thermo scientific, USA). One microgram RNA was shipped to Genewiz (NJ, USA) on dry ice and used for polyA selection, and cDNA library was generated and sequenced using Illumina HiSeq 2×150 bp sequencing. The generated reads were trimmed and aligned to the poplar genome (*P. trichocarpa* v3.1.1) available at Phytozome (https://phytozome.jgi.doe.gov). Based on >80% uniquely mapped reads, we used data from 6 and 3 replicates of AE and its control roots, respectively, and 4 replicates of each AI and its control roots. Finally, differential gene expression analysis was performed using DESEQ2 platform (Love et al. 2014).

### Statistical analysis

For biomass analysis, root image analysis, and C isotope labeling study, contrast analyses between treatments (AI vs. control and AE vs. control) were performed by T-tests with the built-in statistical function in Microsoft Excel. For transcriptomics and metabolomics analyses, AE and AI treatments were performed independently with their own controls (AI vs. control-AI and AE vs. control-AE) for each tissue separately, and analysis of variance tests (ANOVAs) were used for statistical analysis.

Regarding metabolomics data, in general, LC-MS data commonly represents a larger portion of the entire metabolome of the samples, given the metabolite extracts are directly used for analyses. However, in GC-MS analyses, extracts need to be derivatized, and not all compounds are properly volatilized and detected. Our datasets, as expected, did not differ from those statements. The metabolomic fingerprints of our samples resulting from each platform varied remarkably in the number of detected features, with a total of 13,610 features for the LC-MS dataset and 303 features for the GC-MS dataset. Therefore, given that not all the samples could be run in both LC-MS and GC-MS platforms, the datasets derived from each method were not merged into a single dataset, and different statistical analyses were performed on each of the datasets separately.

Multivariate analyses such as permutational multivariate analysis of variance (PERMANOVA) and principal component analysis (PCA) were performed on the LC-MS data, given the larger representation of sample metabolomes. PERMANOVAs were performed with *adonis* function of *vegan* package (Oksanen et al. 2020) to test for significant differences between treatments (AE vs. Control-AE; AI vs. Control-AI) on the overall metabolome structure of the different poplar tissues. In addition, PCAs were used to plot the distribution of study cases of each tissue across the plane formed by the resulting principal components 1 and 2 (PC1 and PC2) and understand the percentage of the total metabolome variability explained by the effect of the treatment (AE or AI). For each PCA resulting from each tissue and treatment, as performed in other plant metabolomics studies (Rivas-Ubach et al. 2017), the score coordinates of the PC axis (PC1 or PC2) which better clustered the treatment (AE or AI) from their control (control-AE or control-AI) were submitted to one-way ANOVA to test for significant clustering differences between AE and control-AE, and AI and control-AI along the PCs. PCAs were performed using both *imputePCA*, to impute missing data, and *PCA* functions of missMDA (Josse and Husson 2016) and FactoMineR (Lê et al. 2008) packages, respectively.

For each tissue, the contrast of treatments (AE and AI) vs. their respective controls (control-AE and Control-AI) of individual metabolomics features, identified and non-identified, one-way ANOVAs were used for all features in both LC-MS and GC-MS datasets. ANOVAs of individual features were performed using *aov* function of *STATS* package in R (R Core team 2021).

In this manuscript, we present significantly changed data (p<0.05) compared to controls; however, in some cases we also include the moderately changed data (0.5<P≤0.1) compared to controls.

## RESULTS

### Polar auxin transport modulators altered root dry weight and branching pattern

To understand the source (leaves) to sink (stem, roots) C allocation in poplar, we used foliar spraying of polar auxin transport modulators, i.e., auxin enhancers (AE) and auxin inhibitors (AI) (Fig. 1), as a tool to increase/decrease the sink allocated C pool and sink strength of growing poplar plants (for convenience, AE or AI-treated plants will be referred to as AE-plants or AI-plants in the manuscript, and the tissues from respective plants will be referred to in a similar manner (e.g., AE-leaf tissue). After 11 days of foliar spraying of AE and AI, the harvested biomass showed no significant changes in leaf and stem dry weights (Fig. 2). However, AE plants showed a moderate increase in root dry weight compared to control (non-treated) plants (27.4% increase in AE vs. control, p=0.13) and a significant increase compared to AI-plants (65.4% increase in AE vs. AI, p=0.01) (Fig. 2 and Supplementary table S1), highlighting the role of AE foliar spraying in increasing root biomass. To further investigate the phenotypic effects of polar auxin transport modulators on sink tissues, we used an image-based root phenotyping approach to evaluate the impact of AE, AI, and control treatments on lateral root development, root length, and maximum root width. Our root phenotyping data showed an increase in lateral root branching frequency in AE-plant roots compared to control roots (frequency: AE=0.36 vs. control =0.13, p=0.05) (Fig. 3 and Supplementary table S2), affirming the role of elevated auxin transport in lateral root formation (increasing sink size). However, changes in the lateral root branching frequency of AI-plants were insignificant compared to controls. Other root system architecture properties, including total root area and width max, were significantly reduced in AI-plant roots compared to the control and AE-plant roots (Fig. 3 and Supplementary table S2). The observed shifts in root growth patterns demonstrated the effectiveness of foliar application of AE and AI in generating a contrasting platform for studying sink tissue growth and source-to-sink communication process in poplar. Following morphological and phenotyping analyses, we performed transcriptome and metabolic profiling of source and sink tissues to identify molecular drivers of sink strength. Based on our experimental design (Fig. 1), the genes and metabolites that were highly expressed or accumulated under AE-condition (vs. controls) but down-regulated/less accumulated or not changed under AI-condition (vs. controls) were considered as “molecular drivers” of sink strength (see below).

**Figure 2.**
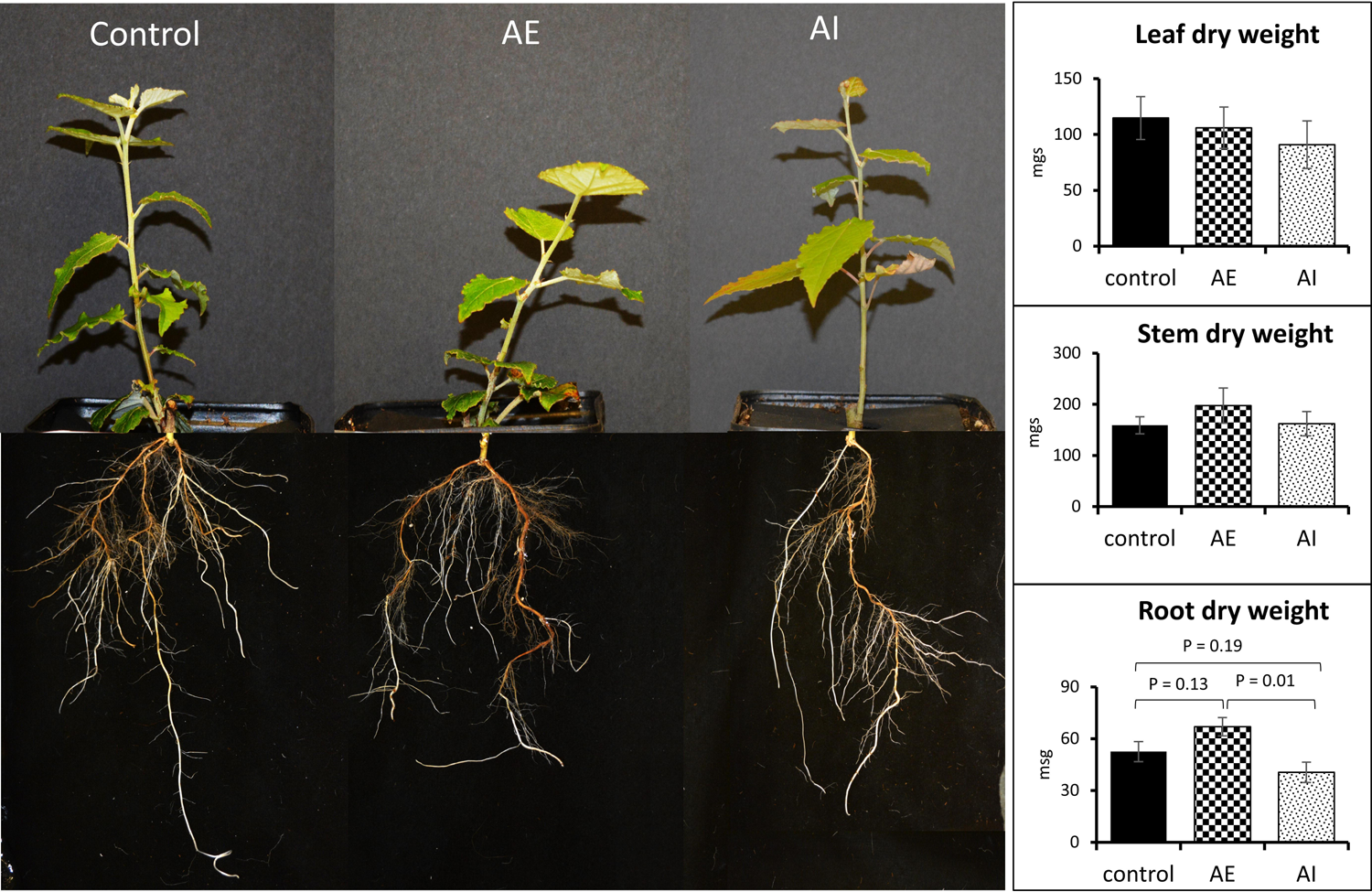
Foliar administration of auxin enhancers (AE) and inhibitor (AI) modulates root dry weight. Representative images from control (n=8), AE (n=6) and AI (n=8) treated biological replicates are shown. Average dry weight measurements calculated from leaf, stem, and root tissues are presented using bar charts. T-test was used for statistical analysis and error bars represent standard error.

**Figure 3.**
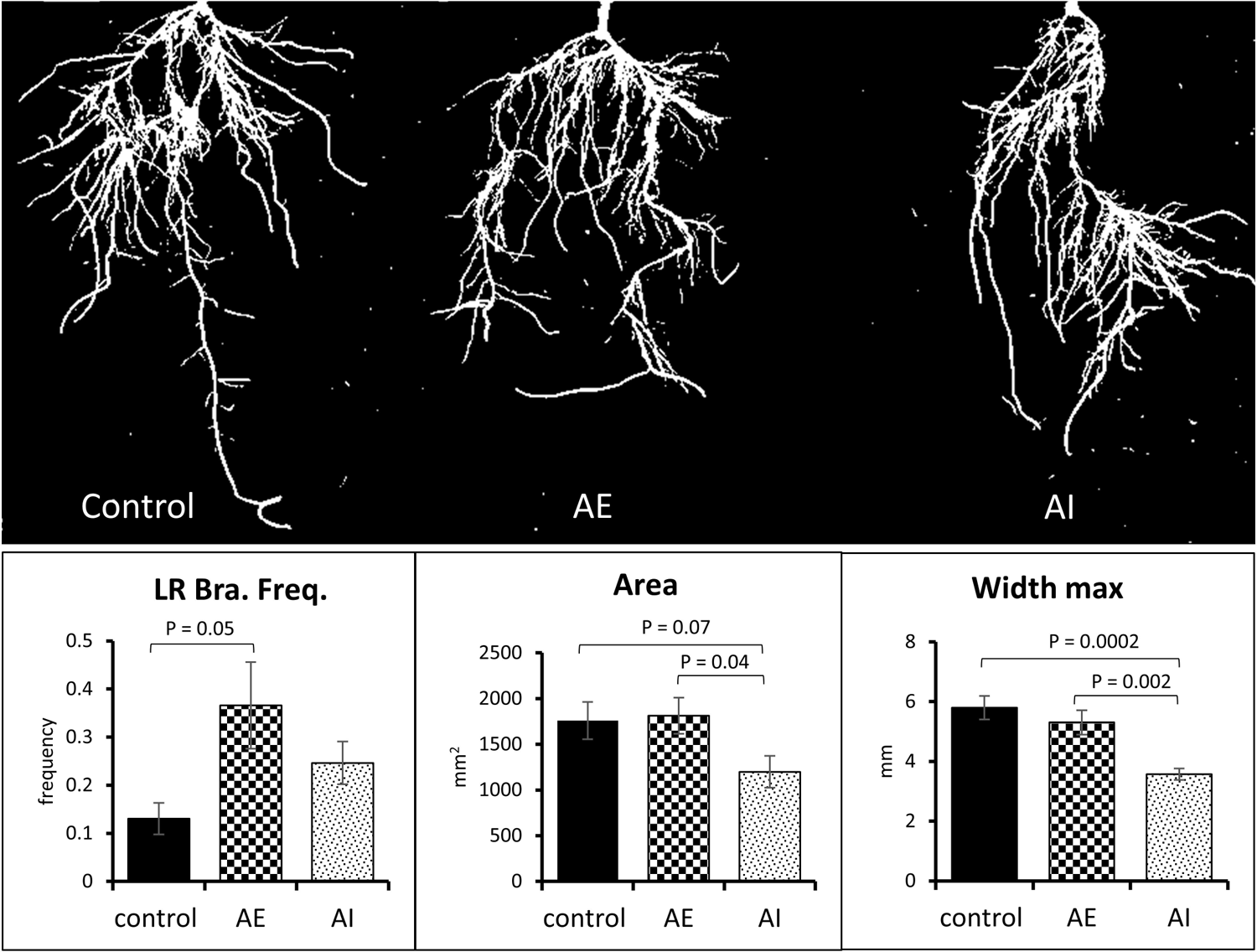
Foliar administration of AE and AI altered root system architecture. The plant root image analyses showed significant differences in root parameters, including lateral root branching frequency (LR Bra. Freq.), root area, and maximum root width (width max). Data averaged from control (n=6), AE (n=7) and AI (n=8) treated biological replicates. T-test was used for statistical analysis and error bars represent standard error. AE: auxin enhancer; AI: auxin inhibitor.

### Transcriptomic analysis of root tissue identified genes involved in sink growth and development

To identify the transcriptional regulators in sink tissue, we performed RNA-seq analysis in the root tissue of hormone-treated plants compared to the control (un-treated) plants. Under AE-treated condition, 42 differentially expressed genes (DEGs) (padj <0.05; Fig. 4) were identified in root tissue, while no significant DEGs were identified in AI-plant (Supplementary table S3).

**Figure 4.**
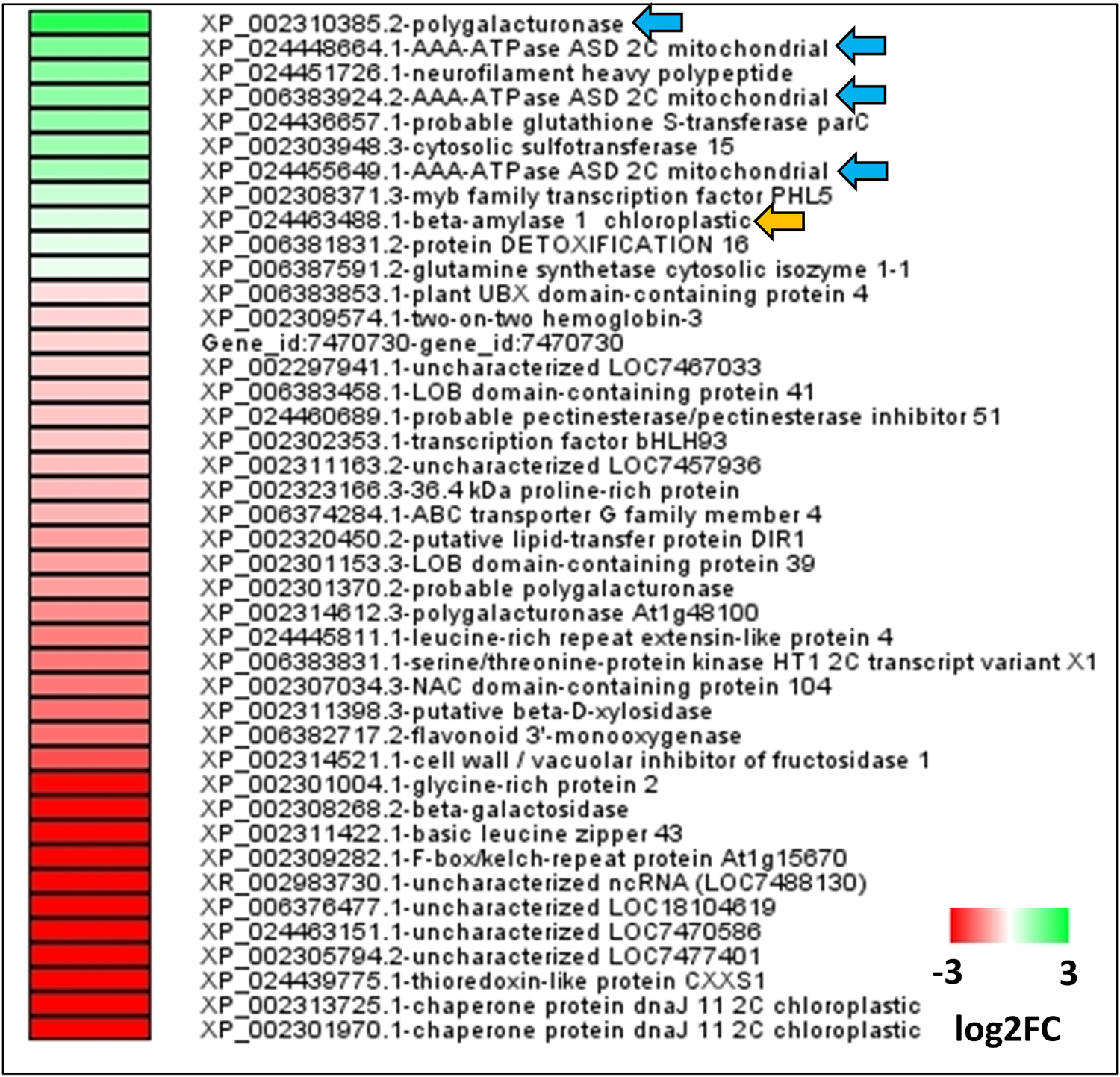
Transcriptomic analysis of AE-root tissue. Heatmap showing the log2 foldchange values of 42 differentially expressed genes (DEGs) in root tissue of AE-treated plants compared to controls. Green color represents upregulated genes, and red color represents down-regulated genes. Arrowheads highlight key upregulated genes involved in plant metabolism that play roles in lateral root growth (blue arrows) and starch catabolism (orange arrow). DESEQ was used for statistical analysis (Padj <0.05), and data was averaged from control (n=6), AE (n=3) and AI (n=4) treated biological replicates. AE: auxin enhancer; AI: auxin inhibitor.

Among 42 AE-root DEGs, 11 were upregulated, and 31 were downregulated (Fig. 4, Supplementary table S3). Among the upregulated genes, the expression of polygalacturonase (Potri.007G144200) (PGA) was 2.5-fold up-regulated under the AE-condition compared to controls. This gene is involved in the cell wall pectin polymer modification process, enabling lateral root outgrowth (Jobert et al. 2021; Peretto et al. 1992). Moreover, three transcripts encoding AAA-ATPases (Potri.004G012700: 1.31-fold, Potri.T104100: 1.56-fold, Potri.004G012500: 1.12-fold) were up-regulated in AE-treated roots. In addition, a transcript encoding a key enzyme in starch degradation, i.e., β-amylase, was up-regulated under AE condition suggesting a possible role of starch metabolism in sink tissues. The upregulation of PGA, AAA-ATPases, and β-amylase genes in roots under AE condition highlights their possible roles in root growth and increasing sink size in poplar, aligning with our earlier observations of significantly increased lateral root branching and a moderate increase in root dry weight under AE condition (Figs 2 & 3).

### Untargeted metabolomic analysis revealed key metabolites as potential molecular drivers of sink strength

The photosynthetically fixed carbohydrates and their metabolism and translocation are critical for sink growth. Therefore, we performed metabolomic analysis of leaf, stem, and root tissues from AE- and AI-plants and their respective controls (AE vs. control-AE and AI vs. control-AI). Permutational multivariate analysis of variance (PERMANOVA) performed on entire metabolome data found significant differences (P <0.05) in the overall metabolome of root tissues under AI-treatment (AI-root vs. control root), and marginal significant differences (P <0.1) in the overall metabolome of stem tissues under AE treatment (AE-stem vs. control stem) (Supplementary table 4A). The overall metabolome of the rest of treatments and tissues did not change significantly compared to their controls (Supplementary table 4A). However, the PCAs (principal component analysis) performed on each tissue showed a significant clustering of scores of hormone-treated (AE and AI) and control samples along the PC1 axis (Supplementary Fig. S1) indicating a degree of difference between the groups. The PCAs showing larger clustering of treatments across the PC1 were in accordance with the PERMANOVA results (Supplementary table 4A).

From all significantly changed features identified from AE-plants, 21, 38, and 38 metabolites matched to specific metabolites in leaf, stem, and root tissues, respectively (ANOVA, p<0.05, supplementary table S4B). Among the significantly altered features in AI-treatment, 32, 38, and 42 metabolites matched to specific metabolites in leaf, stem, and root tissues, respectively (ANOVA, p<0.05, supplementary table S4C). We also included some metabolites that showed a moderate increase or decrease in their relative abundance levels in AE or AI tissues vs control tissues (0.5<P≤0.1) and marked them with single asterisk (*) in figure 5. We could not detect indole-3-acetic acid (IAA) in our samples in any of the tissues or treatments through our untargeted metabolite profiling method. This is probably due to the lower concentration of these hormones in plant tissues that usually require specific sample purification methods followed by targeted approaches using triple quadrupole mass spectrometry (Novák et al. 2012; De Zio et al. 2019).

**Figure 5.**
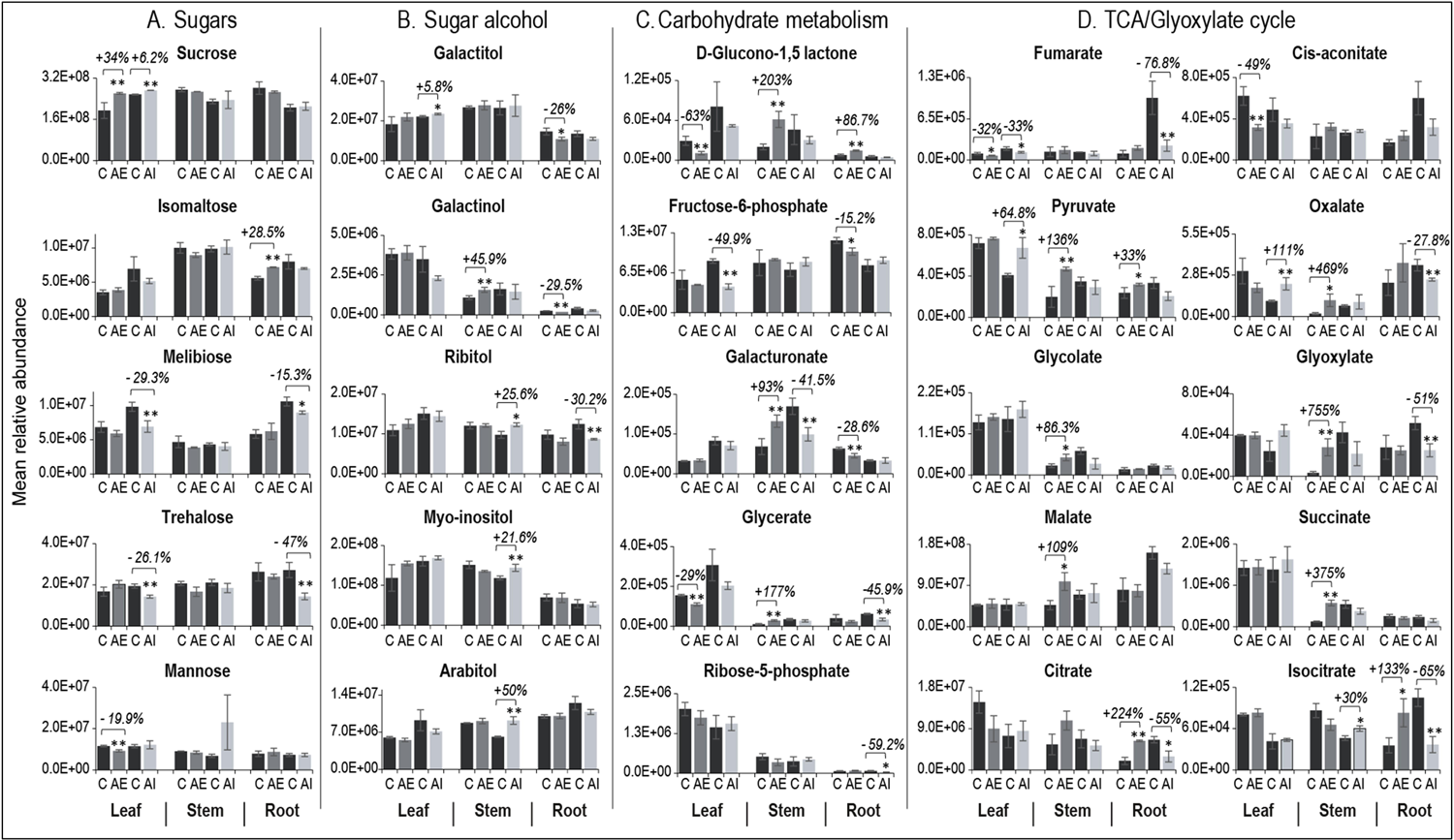
Metabolomic profiling of selected metabolites in leaf, stem, and root tissues of AE and AI foliar-sprayed plants compared to their controls. Relative abundance levels of **(A)** sugars, **(B)** sugar alcohols, **(C)** carbohydrate metabolism (glycolysis, pentose phosphate pathway and other pathways), and **(D)** TCA/Glyoxylate cycle intermediates are presented. The percentage change in relative abundance levels of significantly altered metabolites in AE vs C and AI vs C are presented in % values above the bars. Significantly altered metabolites between treated plants (AE or AI) and their respective controls are shown with ** for P < 0.05 and with * for 0.05<P≤0.1. One way ANOVA was used for statistical analysis, and data averaged from n= 3-6 biological replicates with 3 technical replicates. AE: auxin enhancer; AI: auxin inhibitor.

Further, we compared the overall profile of identified metabolites in source vs. sink tissues between treatments to see if relatively more metabolites with higher relative abundances were found under AE-treatment, especially in sink tissues. Interestingly, in source leaf, AI-treatment resulted in more metabolites with higher relative abundances (50% of identified metabolites had higher abundance levels in AI-leaf vs. 33.3% in AE-leaf), while in sink tissues, including stem (36.8% in AI-stem vs. 76.3% in AE-stem) and root (19% in AI-root vs. 23.6% in AE-root), AE-treatments resulted in more metabolites with higher relative abundances (Supplementary Fig. S2 and supplementary tables S4B & S4C). This indicates that AE induced a higher relative metabolic activity in sink tissues while AI induced higher relative metabolic activity in source leaves.

Metabolites with significantly altered abundances between experimental treatments were categorized into seven broader KEGG biological pathway categories. The total number of significant metabolites in each pathway category (Supplementary Fig. S3A), and relative higher or lower fold change in abundance levels were plotted (Supplementary Fig. S3B). We also developed a comprehensive view of the adapted KEGG metabolic map for each tissue under all conditions highlighting the metabolites with higher or lower foldchange in abundance (Supplementary Figs. S4A, S4B, S4C). More specifically, we plotted a selective set of the potential metabolites associated with source-sink C relationships and sink strength including sugars, sugar alcohols, carbohydrate metabolism intermediates, and TCA/Glyoxylate cycle intermediates (Fig. 5A,5B,5C,5D) which we describe in more details in the following sections. From here onward, the % metabolite level refers to the % relative abundance level of a specific metabolite in hormone-treated tissues in reference to the same metabolite’s abundance levels detected in controls.

### Sugar alcohols and products of starch metabolism are associated with sink activity in poplar

The non-photosynthetic sink tissues, such as root, demand mobile forms of C, e.g., sucrose and sugar alcohols, from the source leaves for sink establishment and maintenance. Therefore, we assessed the relative abundance levels of these C forms in hormone-sprayed plant tissues. Through foliar spraying of AE and AI, a 34% and 6.2% increase in sucrose levels were observed in source leaves, respectively, while sucrose levels in sink stem and root tissues remained unchanged compared to controls (Fig. 5A). Apart from sucrose, sugar alcohols such as galactinol act as mobile C forms in several tree species (Dominguez and Niittylä 2021). Therefore, we looked at the sugar alcohol levels in stem and root tissues. We observed a 45.9% increase in galactinol levels in stems under AE-condition (Fig. 5B). Moreover, spraying with the auxin transport inhibitor increased the levels of three other sugar alcohols, including arabitol (+50%), ribitol (+25.6%), and myo-inositol (+21.6%) in the AI-stem (Fig. 5B). In root tissue, we found reduced levels of galactitol (−26%) and galactinol (−29.5%) (two major source-to-sink mobile C forms) under AE condition (Fig. 5B), highlighting a possible important role of sugar alcohols in sink tissues. Furthermore, focusing on sugar metabolism in roots, we found a 28.5% increase in the levels of sugar isomaltose (a starch breakdown product) under AE condition (Fig. 5A), signifying the role of starch metabolism in sink tissues. On the other hand, the AI-root tissue showed reduced levels of two other sugars, melibiose (−15.3%) (a breakdown product of raffinose) and D-trehalose (−47%) (involved in sugar sensing and starch metabolism) (Fig. 5A). Overall, we found multiple mobile sugar alcohols and starch metabolism-derived sugars as potential molecular drivers of sink growth and activity in poplar.

### TCA cycle metabolites showed an opposite trend in root tissue under AE and AI conditions

The mobile C forms translocated from source tissues are metabolized in sink tissues via cellular respiration processes that comprise glycolysis, pyruvate oxidation, and TCA/Glyoxylate cycle, in which the TCA cycle yields NADH for downstream ATP synthesis, while the glyoxylate cycle replenishes the TCA cycle intermediates for sugar biosynthesis. Under AE condition, source leaves showed no significant changes in metabolites involved in glycolysis and pyruvate metabolism while they did show reduced levels of TCA cycle metabolites fumarate (−32.2%) and cis-aconitate (−49%)) (Fig. 5C). In contrast, during AI conditions, reduced levels of β-D-fructose-6-phosphate (−49.9 %) and fumarate (−32.8%), and higher levels of pyruvate (+64.8%) and oxalate (+111.4%) were found in the leaves (Fig. 5C & 5D), indicating a shift in cellular respiration activity in source leaves.

In the stem tissue, very high levels of pyruvate (+135%), glycolate (+86%), glyoxylate (+755%), malate (+109.4%), oxalate (469.6%), and succinate (+375%) (Fig. 5C) were observed under AE condition, which shows a very high rate of cellular respiration in this sink tissue. On the contrary, under the AI condition, the stem tissue showed no changes in glycolysis- and pyruvate-associated metabolites, while higher levels of isocitrate (30%) were detected (Fig. 5C).

In the sink root tissue, AE condition resulted in lower levels of a glycolysis-related metabolite, β-D-fructose-6-P (−15.2%), while citrate levels showed a significant higher accumulation (up to 224%) (Fig. 5C & 5D). However, under AI condition, four TCA/glyoxylate cycle intermediates (citrate (−55%), isocitrate (−65%), glyoxylate (−51%), fumarate (−76%)) and two other related metabolites (oxalate (−27%), itaconate (−84%)) were significantly reduced in abundance (Fig. 5C). The observed opposite trends under AE and AI conditions clearly suggests a very important role of TCA cycle intermediates and specifically citrate, as a key molecular driver of sink strength, to meet energy demands during sink development.

### Auxin transport modulators alter metabolite abundances in diverse metabolic pathways in sink tissues

In addition to sugar alcohols and respiration-associated metabolites, physiological and developmental processes in sink tissues require other C intermediates, including pentose sugars generated via the pentose phosphate pathway (PPP), secondary metabolites, phytohormones, nucleic acids, and amino acids. In both AE and AI conditions, source leaf tissue did not show any significantly induced PPP metabolites. However, in the sink stem tissue, AE-treated plants showed higher levels of D-glucono 1,5 lactone (+203%), and glycerate (+177.6%) that participate in PPP, while their levels remained unchanged in AI-plants (Fig.5D and Supplementary tables S4B & S4C). Interestingly, in sink roots, only AE-plants exhibited higher levels of D-glucono 1,5 lactone (+86.7%), indicating a possible role of PPP intermediates to support sink growth under AE condition. In association with PPP, we found elevated levels of nucleic acids such as adenosine and 5,6 dihydro uracil in AE-roots suggesting their possible roles in supporting cell division and elongation processes in root tissue. On the other hand, the AI-treated roots showed a lower level of non-oxidative PPP intermediate, Ribose-5-phosphate (−59.2%) (Fig.5D and Supplementary tables S4B & S4C).

Another notable difference was found in the levels of galacturonic acid, which compose pectin polymer of growing cell walls. We observed a higher level of galacturonic acid (93%) in AE-stem tissues, while we noticed an opposite trend in AI-treated stems (−41.5%) (Fig. 5D). Interestingly, in sink roots, the AE-plants showed reduced levels of galacturonic acid (−28.6%), thereby highlighting its potential involvements in sink tissue activities possibly through providing C pools.

We found a very active secondary metabolism in the sink stem tissue of AE-plants demonstrating higher relative abundance levels of 14 out of 17 significantly identified secondary metabolites (Supplementary Fig. S4B and Supplementary tables S4B & S4C). Among them, fisetin was found at a very high level (+960%), reported to play a role in stimulating IAA transmembrane transport and blocks NPA binding (Faulkner and Rubery 1992), highlighting the activation of metabolites in support of polar auxin transport in AE-stem. Other notable secondary metabolites of higher abundance in sink stem tissue include 4-hydroxy benzoic acid (+1441%) and chlorogenic acid (+1398%), which participate in cell wall phenolic polymer biosynthesis. Chlorogenic acid was also found at a higher abundance level (+169%) in the root tissue of AE-plants (Supplementary tables S4B & S4C, and Figs S4A, S4B, S4C). On the other hand, the AI-treated source leaves showed higher levels of flavonoids such as naringin (only found in the AI-leaf and not detected in the control leaf) and ferulate (+96%), among which a non-glycosylated form of naringin (naringenin) was reported to negatively regulate polar auxin transport process (Peer et al. 2004; Brown et al. 2001; Peer and Murphy 2007; Rasouli et al. 2016). In AI-stem and root tissues, most secondary metabolites were reduced in their abundance, out of which the level of fisetin was reduced (−99%) in root tissue, which again could signify its role in influencing auxin transport to the roots in poplar. Overall, the secondary metabolites identified in source and sink tissues could play a role not only in sink growth and activities but also in regulating polar auxin transport in poplar.

Regarding hormone metabolism, even though phytohormones are not easily detectable with untargeted metabolomic methods, we obtained signals that matched against jasmonic acid and gibberellins m/z and RT (retention time) values, which showed interesting trends in the sink tissues. Under AE condition, gibberellin was elevated in abundance in sink stem tissue (only found in the AE-treated stem and not detected in controls), while the level of this phytohormone was reduced in the stems of AI-plants (−48%). It was shown that auxin transport inhibitor NPA reduced gibberellin levels in plants (Ross 1998). Gibberellins were also found to enhance sink strength (Roopendra et al. 2018) and regulate sugar alcohol levels (Zhuang et al. 2015). The level of jasmonic acid was not detected under the AI-treated conditions in stems, while it was reduced in the root tissue (−81.4%) (supplementary tables S4B & S4C and Figs S4A, S4B, S4C).

We also looked at the metabolism of amino acid and nitrogenous compounds in source and sink tissues of hormone-treated plants. In AE-plants, the level of amino acid phenylalanine (+214%) was elevated in abundance in source leaves, while in stem tissue higher levels of tyramine (a breakdown product of tyrosine) (+52.8%) were detected. Both amino acids act as a precursor in secondary metabolite biosynthesis in plants (Feduraev et al. 2020), supporting a higher level of secondary metabolism found in AE-stem tissues (supplementary tables S4B & S4C and supplementary Figs S4A, S4B). AE-treated stems accumulated a higher level of amino acid alanine (+231%), which is interconvertible with pyruvate (Diab and Limami 2016). On the contrary, in sink tissue, the levels of all amino acids were significantly reduced, including notable amino acids glutamine (−35%), proline (−34%), and L-alanine (−26%) (supplementary tables S4B & S4C and Fig. S4C). Under AI-condition, the glutamate-derived GABA (Gamma-aminobutyric acid), which was found to play a role in carbon-nitrogen metabolism and also in regulating C fluxes into the TCA cycle (Bouché and Fromm 2004), was only found in AI-plant roots and not detected in controls. Moreover, the amino acid serine showed higher accumulations in AI-roots (+24.15%) while showed a significant reduction under AE condition (−22.8%), highlighting a potential role of serine in root growth and maintenance (supplementary tables S4B & S4C, and Figs S4C).

### AE-plants maintained a higher net fixed C export from source to sink tissues

The physiological, transcriptomics and metabolomic analyses revealed that foliar spraying of AE significantly increased root branching frequency and moderately increased root dry weight (27.4% increase in AE vs. control, p=0.1) likely facilitated by metabolic drivers such as shifts in the abundances of sugars, sugar alcohols, carbohydrate metabolism products, and TCA cycle intermediates (Figs. 2-5). To further evaluate whether these phenotypic and molecular changes were primarily due to photosynthate transfer from source to sink tissues, we performed a ^13^CO_2_ tracer experiment. Based on our phenotypic and molecular data, we expected to see a higher enrichment of ^13^C in sink tissues under AE condition indicating the mobilization of the photosynthetically fixed C from source leaves to sink root tissues.

Our results showed the highest δ^13^C values within photosynthetic tissues (i.e., leaves), with both treatments and controls having nearly identical δ^13^C values (∼ 4000‰) (Fig. 6, supplementary table S5). The sink tissues (stem and root) were segmented to evaluate if there were any spatial differences in the ^13^C isotope enrichment levels. Stem-base (2-5cms region of the stem tissue close to the roots) is very critical in establishing and maintaining adventitious (shoot-born) roots and shown to accumulate precursor carbohydrates, hormones, and other regulatory genes required for root formation and growth (Agulló-Antón et al. 2014, Druege et al. 2019). Therefore, we looked at ^13^C enrichment levels in the stem-base and stem-top (region without stem-base), separately (Fig. 6). The migration of the fixed ^13^C signal into stem segments did not change significantly in AE-plants, while AI-treated plants showed a moderate reduction in ^13^C enrichment levels in their stem-base (∼278 ‰) compared to controls (∼504 ‰, P=0.16) and AE-plants (∼628 ‰, P=0.21) (Fig. 6).

**Figure 6.**
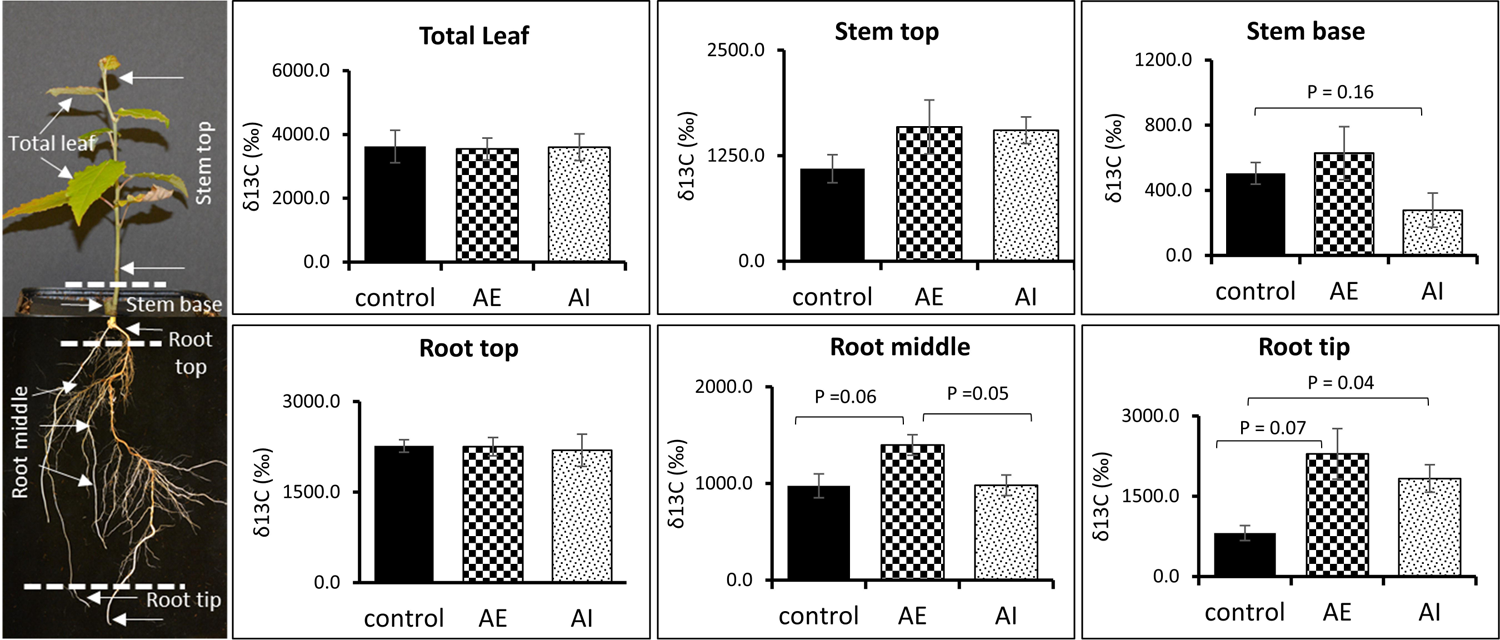
Foliar administration of AE and AI altered the allocation of source leaf fixed ^13^CO_2_ to the sink tissues. 11-days post-hormone spraying, the plants were moved inside a Plexiglass chamber enriched with ^13^CO_2_ for 4-days. The isotope enrichment levels in different tissues and root segments measured by isotope ratio mass spectrometer (IRMS) are presented. Data averaged from control (n=4), AE (n=4) and AI (n=4) treated biological replicates with 3 technical replicates. T-test was used for statistical analysis, and error bars represent standard error. AE: auxin enhancer; AI: auxin inhibitor.

To evaluate how effectively our AE- and AI-treatments modulate C transport into the root tissue, we specifically screened the C assimilation levels in segments of root tissue that include root-top (root region close to stem base), root-middle (a region that comprises most of the main roots with a higher frequency of lateral roots), and root-tip. The root-top tissue showed similar δ^13^C levels under both treatments and control conditions (∼2000-2300 ‰). Further, the AE-plants showed a moderate increase in ^13^C assimilation levels in the root-middle region (∼1400 ‰) compared to the control (∼975 ‰, P=0.06) (Fig. 6), signifying that AE-treatment led to increased deposition of photosynthetically derived C into sink roots, especially in the region of lateral root growth and emergence, which can be corroborated with the higher lateral root branching frequency observed in AE-plants (Fig. 3). On the other hand, AI-plants maintained a similar level of ^13^C assimilation in their root-middle region compared to control plants (Fig. 6). In actively dividing root-tip region, interestingly a higher δ^13^C level was found in both AE-(∼2290 ‰, P=0.07) and AI-root tips (∼1830 ‰, P=0.04) compared to controls (∼812 ‰). However, it should be noted that the root tip routinely synthesizes auxins locally (Ljung et al. 2005), and further studies are required to understand the direct effects of foliar sprayed AE and AI on root tip synthesized auxin levels Thus, the stable isotope tracer experiment is consistent with our earlier observations of increased root branching frequencies, a moderate increase in root dry weight, and altered primary carbohydrate metabolism in AE-plants (Figures 2 and 3). These results suggest an elevated demand for C pools and increased sink strength in root tissues in plants treated with auxin enhancers.

## Discussion

The process of C allocation is a function of source-sink interactions (Coleman and Isebrands 1994). However, the main control of C allocation between source and sink tissues is suggested to lie in sink size and activity (Ho 1988). Therefore, factors affecting sink strength are possibly the most important for regulating C allocation and partitioning (Gifford and Evans 1981). In this work, we used auxin modulators as chemical tools to alter sink strength and to understand C dynamics in growing poplar plants. We integrated biomass measurements, root architectural phenes, ^13^C assimilation, metabolite profiles, and gene expression levels to understand the molecular regulation of sink strength in poplar. Although both stem and root tissues are considered as major sink organs in woody plants, this research is mainly focused on the root tissue and the roles of primary carbohydrate metabolism in root growth and physiology under auxin-mediated altered C sink condition.

Auxins are primary growth regulators that modulate almost every aspect of plant physiology, growth, and development. In this study, we amplified and attenuated the polar auxin transport from source-to-sink tissues through foliar (leaf) spraying of AE and AI. The positive effects of foliar applications of AE on plant growth include increased root/shoot length and dry mass as previously reported in *Brassica juncea* (Mir et al. 2020) *Vigna radiata* (Ali et al. 2008), *Petunia hybrida* (Ahkami et al. 2013), *Solanum melongena* (Hayat et al. 2006) and soybean (Sarkar et al. 2002). Opposite trends have been reported in the case of AI applications (e.g., NPA) in other plant species, including decreased taproot growth in spruce (Rincón et al. 2003). These observations could be linked to auxin’s roles in stimulating cell division, cell elongation, and cell differentiation (Sitbon and Perrot-Rechenmann 1997). Moreover, the stimulatory effect of auxin on plant growth and development is also associated with its impact on higher production of the carbohydrate content in plants (Ahkami et al. 2013; Sh and Dawood 2013). Previous studies showed the stimulatory effects of auxin application in the mobilization of carbohydrates in source leaves, elevated the translocation of assimilates, and enhanced sugar availability in sink (root) tissues (Haissig 1974; Altman and Wareing 1975; Haissig 1986; Husen and Pal 2007; Agulló-Antón et al. 2011). Furthermore, Mishra et al. 2009, observed that auxin homeostasis and related control of sink tissue development can be modulated by sugar signaling (Mishra et al. 2009). Therefore, in our study, we hypothesized that enhancement or repression of phloem-transported auxin levels could modulate C allocation and potentially reveal the molecular drivers responsible for long-distance source (shoot)-sink (root) C relationships and sink strength in poplar.

In our auxin-mediated sink strength modulation experiment, the AE-plants showed a moderate increase in root dry weight (Fig. 2) over the control plants, while the AI-plants showed a moderate reduction in root dry weight versus the control (Fig. 2), thereby creating contrasting platforms to study the metabolism involved in C allocation and sink (root) strength in poplar. The increase in root dry weight in AE-plants, without changes in source leaf dry weight or leaf counts in AE-plants vs. controls (Fig.2, Supplementary table S1), indicates that AE foliar spraying modulated exclusively sink strength in poplar. Further, we performed image-based root phenotyping to screen specific root architectural parameters under the two contrasting sink conditions and observed an increase in lateral root branching frequency upon treatment with AE (Fig. 3). In contrast, AI-plants showed no significant changes in lateral root branching frequency but displayed reduced root area and maximum root width compared to controls (Fig. 3). These results highlight that the increase in root dry weight in AE-plants could be primarily driven by increased lateral root density, while a reduction in root dry weight in AI plants could be mainly governed by decreased root area and root thickness-related traits. In AE treated plants, we did not observe a correlation between lateral root branching frequency and the total root area, where the lateral root branching frequency increased while the total root area showed no changes compared to controls (Fig. 3). To capture all the thin lateral roots by imaging, we placed the root system in a black tray with water to evenly spread all lateral roots. However, during image processing and trait extraction using the DIRT software package, we noticed that increasing the threshold to capture all lateral roots could lead to increasing background noise in other root traits. So, one of the possible reasons for observing just minor effects on root area could be the technical limitations associated with thresholding-based root image analysis. Other possible reasons for not observing a more pronounced root phenotype in AE- and AI-treated plants versus controls include i) the concentrations of AE and AI hormones used for foliar spraying (see below), ii) directing the synthetic auxins toward root tissue primarily via source leaf absorption to mimic naturally enhanced/repressed polar auxin flow (compared to studies that directly apply auxin modulators on growing roots), which could limit the optimal on-site auxin concentration to significantly impact root growth, and iii) the root developmental stage in our study (well-established 4-5 weeks old root system prior to foliar spraying) in which the sprayed auxin modulators mainly impacted lateral root growth rather than establishing newer (adventitious) roots. Although foliar application of phytohormones, fertilizers, and micro and macronutrients is known to enhance plant growth by improving photosynthesis and resource allocation, the cuticular penetration of sprayed components are considered as rate-limiting factors in foliar absorption (Kerstiens 1996). In future studies, optimizing a more efficient and precise delivery of hormones and their concentrations could increase a higher sink strength level, thereby obtaining more pronounced and measurable root phenotypes for image-based studies.

Sink establishment and maintenance are facilitated by a continuous biosynthesis, deposition, and modification of cell wall polymers that comprise the structural carbohydrates of plants. Auxin regulates cell expansion and cell wall biosynthesis through cell wall modification genes including polygalacturonase (Jobert et al. 2021; Peretto et al. 1992), expansins, xyloglucan endotransglucosylase (XTH) (Majda and Robert 2018). Polygalacturonase is a pectin remodifying enzyme exclusively expressed under auxin’s influence to loosen the cell walls to facilitate the lateral root emergence process (Jobert et al. 2021; Peretto et al. 1992). Interestingly, the gene expression analysis of AE-roots identified a transcript encoding polygalacturonase (Fig.4, supplementary table S3), reaffirming an active lateral root (sink) growth under AE condition as highlighted by our root phenotypic results (Figs 2 and 3). This candidate gene combined with other lateral root growth-related genes induced in AE-roots (marked by blue arrows in Fig. 4 and listed in supplementary table S3) stand as promising candidates to determine their functions in increasing sink strength in poplar and other related tree species.

Metabolomic analysis revealed differences in leaf, stem, and root tissues between AE and AI-treatments compared to controls (Fig. S1; PC1 axis of PCA plots), indicating a shift in metabolism as a result of our auxin-stimulated carbon sink approach which could explain the observed phenotypic differences in root tissue (Figs. 2 and 3). To assess whether these metabolic shifts in sink tissues can be associated with source-to-sink C allocation, we performed ^13^CO_2_ enrichment of hormone-sprayed plants and control plants to evaluate whether the treated plants maintained a higher/lower C allocation from source leaf to sink tissues compared to controls. While AE-plants retained a moderate increase in ^13^C allocation to stem segments, *i.e.,* stem-base and stem-top, such as, they showed a significantly higher C allocation to the root region with a high frequency of lateral roots (root-middle) compared to controls (Fig. 6). While we observed a higher allocation of ^13^C to root middle region, indicating a possible increase in sink strength in the form of lateral roots, AE plant roots did not show an increase in total root area. Therefore, another possible scenario could be redistribution of assimilates between root segments toward branching (Sharkey 2015). However, the AI-plants demonstrated similar ^13^C enrichment levels in leaves, stems, and root-middle region compared to control plants (Fig. 6). This later observation is likely attributable to the relatively lower AI concentration used in this study as foliar application (100 μM NPA). Other studies in woody plants, for example in Oak, used a higher concentration of up to 200 μM of NPA (Valladares et al. 2020), which was directly applied at the site of root initiation to inhibit root formation. Despite Auxin being a plant growth regulator, a higher concentration of foliar sprayed auxin negatively impacted plant height and biomass yield (Nassef and El-aref 2018; Di Benedetto et al. 2015; Rastogi et al. 2013), therefore highlighting the need to optimize the hormone concentration based on the plant species studied. We tested and used an optimal concentration of AI for foliar spraying to avoid chemical stress on source leaves, as shown by no reduction in leaf number or leaf dry weight under AI-condition (Fig.2 and supplementary file S1). In addition, our C isotope enrichment experiment was conducted for a period of 4 days after completing our spraying experiments, and it is possible that during this 4-day period, AI-plants lacked a higher concentration of AI in foliar leaf tissue to more effectively repress the polar auxin transport, thereby resuming the C allocation rates similar to the control plants. However, we suggest that the sink phenotype (root biomass and associated metabolism) obtained through AE-treatment could primarily be due to increased C flow to sink tissues that altered root metabolism and growth. Therefore, the specific metabolites that were significantly altered in hormone-treated tissues compared to controls can potentially involve in source-to-sink C communication and contribute to sink strength in poplar. Here, we mainly discus the results pertaining to primary carbohydrates, sugar alcohols, starch breakdown products, and TCA/glyoxylate cycle metabolites (Fig. 5 and 7).

**Figure 7.**
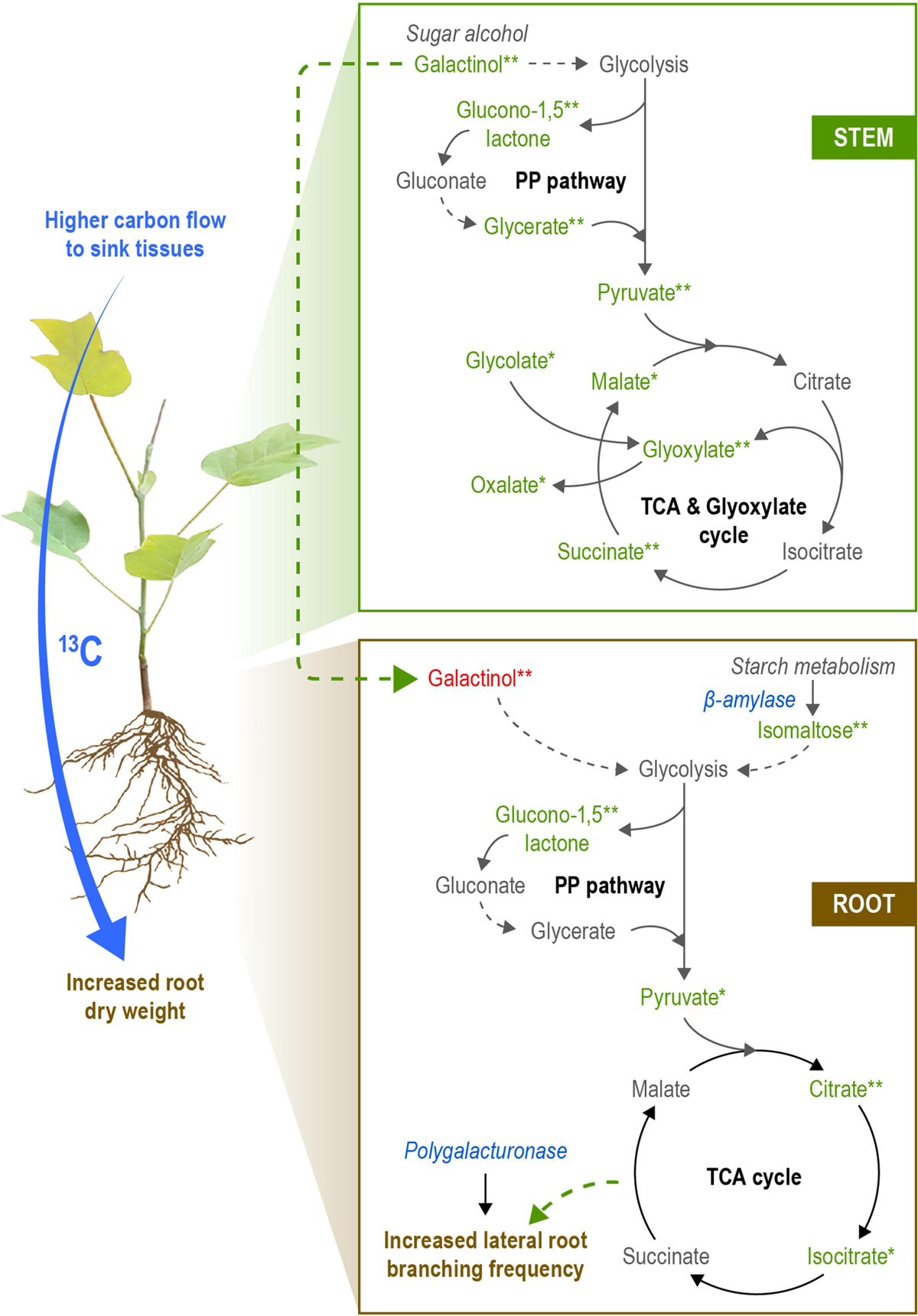
A model depicting source-sink carbon relationships including molecular drivers of sink strength in poplar. We suggest that the source-sink carbon relationships in poplar could be fueled by mobile sugar alcohols and starch metabolism-derived sugars as key molecular drivers of sink strength. We also postulate a very important role of TCA cycle intermediates in providing the energy demands during sink (root) growth. Black solid arrows indicate direct relationships between intermediates derived from the KEGG pathway, while black dashed arrows represent indirect relationships. Metabolites in green and red colors denote an increase and decrease in abundance, respectively (under AE condition). Transcripts in blue color are increased in expression levels under the influence of auxin enhancers. Green dashed arrows highlight our hypotheses on the possibility of galactinol transport from stem to root and the possible role of TCA cycle intermediates in providing energy for lateral root branching and improving sink strength. (** denotes p<0.05 and * denotes 0.05<p≤0.1). PP pathway – Pentose phosphate pathway.

Although sucrose is the major photosynthetically fixed phloem translocated sugar in most plant species, we did not find any significant changes in sucrose levels in sink tissues (root and stem) under both AE and AI treatments. However, we found significant shifts in the levels of different types of mobile sugar alcohols in stem and root tissues (Fig. 5). For example, the stem tissue of AE-plants showed higher levels of the mobile sugar alcohol galactinol (Fig. 5) which is a precursor of raffinose family oligosaccharide biosynthesis in plants. In poplar, a higher phloem galactinol level was achieved by overexpressing the galactinol synthase gene, which led to an increase in phloem sucrose and starch level and starch deposition in xylem tissue (Unda et al. 2017), highlighting the role of galactinol in C metabolism and transport in poplar trees. In another study (Coleman et al. 2007), an attempt to alter C metabolism in hybrid poplar (*Populus alba × Populus grandidentata*) by overexpressing Acetobacter xylinum UDP-glucose pyrophosphorylase led to an increase in sugar alcohols comprising galactinol, galactitol, and myo-inositol which resulted in a higher starch and cellulose levels, underscoring the link between sugar alcohol accumulations and primary carbohydrate metabolism. Also, disruption of sink C allocation in aspen (*Populus tremula*) by stem girdling increased galactinol levels in stem tissue, suggesting a role of this sugar alcohol in C allocation in *Populus sp.* (Lihavainen et al. 2021). In our study, a higher galactinol level in AE-stem tissue indicates a higher C allocation to the stem tissue, which was commensurate with a higher overall trend (but not significant) in ^13^C enrichment levels in stem segments under the same condition (Fig. 6). Significant higher accumulations of glyoxylate and glycerate in stem tissue (Fig. 5) is another indication of increased C pool and metabolism under AE condition. Focusing on root tissue, both AE- and AI-plants showed reduced levels of sugar alcohols in which AE-plants contained lower levels of major sugar alcohols galactinol and galactitol, while AI-plants showed a lower level of ribitol compared to controls (Fig. 5). A lower level of major sugar alcohols in roots (including galactinol) under AE condition indicates either there was a lower translocation of sugar alcohols from the stem (which showed a higher galactinol level) to the root, or consumption of these sugar alcohols during root growth (root branching). Altogether, modulating auxin transport altered the levels of major sugar alcohols, especially in sink tissues, highlighting their possible roles in regulating sink strength in poplar.

Starch content and mobilization play a critical role in C allocation, sink establishment, and sink strength (MacNeill et al. 2017; Hartmann and Trumbore 2016; Noronha et al. 2018; Regier et al. 2010; Song et al. 2019). We found a higher level of isomaltose (derived from the starch breakdown process) (Fig. 6) together with a higher expression level of a beta-amylase transcript in root tissue only in AE-plants (Fig.4), highlighting an increase in the starch breakdown process under AE condition in roots. In Arabidopsis, loss of function of the beta-amylase gene resulted in decreased starch metabolism, which reduced primary root length (Song et al. 2019). Here, the increase in starch catabolism product can be associated with observed higher root biomass and lateral root branching frequency (sink size) (Figs. 2,3). On the other hand, the AI-plants showed reduced levels of D-trehalose in leaf and root tissues (Fig. 5). Trehalose was found to increase starch biosynthesis in Arabidopsis and to induce the expression of beta-amylase gene expression (Wingler et al. 2000), altogether indicating a possible lower starch catabolism in AI-root tissue. The higher level of isomaltose in AE-roots could likely fuel glycolysis and cellular respiration via the TCA cycle, thereby generating ATP and C compounds required for sink activities. However, to validate this assumption, the actual starch content in source and sink tissues should be quantified in future works.

Cellular respiration involves controlled breakdown of sugars that arise from sucrose and starch metabolism through glycolytic pathways to ultimately yield ATP, NADH, and C skeleton via TCA/glyoxylate cycle to support basic cellular activities. In aspen (*Populus tremula*), a reduced sink C allocation by stem girdling approach led to a higher C level in the stem tissue that altered the metabolic flux through TCA cycle metabolites, highlighting the role of the TCA cycle in maintaining carbon-nitrogen balance in poplars (Lihavainen et al. 2021). Plant hormone auxin was found to regulate cellular respiration (Leonova et al. 1985) and in turn, perturbation of TCA cycle metabolism affected the polar auxin transport process (Ohbayashi et al. 2019). Further, auxin-sensitive mutant plants (with altered auxin signaling) were shown to have impairments in the usage of starch that were most likely linked with changes in the levels of TCA cycle intermediates, signifying the relationship between cellular respiration and auxin-mediated growth (Batista-Silva et al. 2019). However, our understanding about the impacts of auxin availability and C pool and their relationships on TCA cycle activation is fragmentary. In sink stem tissue, AE-plants showed a very high level of pyruvate and TCA cycle metabolites, including succinate, glyoxylate, and its precursor glycolate (Fig. 5), thereby suggesting a higher primary carbohydrate metabolism to yield ATP via the TCA cycle and a possible activated sugar biosynthesis via the glyoxylate cycle to support the sink demand. However, in sink root tissue, a significantly higher level of citrate was accumulated under AE condition (while opposite trend observed under AI condition) indicating an increase in TCA cycle activity in roots which could be possibly fueled by the elevated isomaltose levels (Fig. 5, 7) arising out of the starch breakdown process. In poplar, during spring, a transition from bud break to expanding leaves (growing young leaf act as sink tissue) resulted in a peak abundance of starch breakdown product (maltose) accompanied by an increase in TCA cycle metabolites (malate, succinate, 2-oxoglutarate), highlighting the involvement of starch breakdown process and TCA cycle in sink growth (Watanabe et al. 2018). In a recent study, manipulation of TCA cycle-associated genes in Arabidopsis showed a reduction in the starch level and its degradation in leaf tissue which was reported to be associated with a reduced TCA cycle metabolite level, highlighting the link between mitochondrial respiration and starch metabolism (Zhang et al. 2021). Moreover, the mitochondrial respiration process was involved in secondary cell wall formation in roots (van der Merwe et al. 2010), and down-regulation of a TCA cycle gene (mitochondrial malate dehydrogenase) resulted in reduced root area, root weight, and respiration rate in tomato (van der Merwe et al. 2009). Interestingly, in our study, AI-plant roots showed reduced levels of most of the TCA cycle metabolites and glyoxylate levels (Fig. 5), which can be correlated with the observed reduced root biomass and root phenotypes such as root area and maximum root width (Figs. 2,3). Overall, our work highlights an important role of the TCA cycle, and specifically citrate, in enabling plants to import and assimilate C to sink tissues under an auxin-modulated C-enriched condition in poplar (Fig. 7).

In summary, this is the first comprehensive study of source-sink C relationships in trees like poplar where foliar treatments of auxins were applied to modulate carbon allocation from source leaf to sink root tissues. Through foliar spraying of synthetic auxins, we developed a novel contrasting C allocation/demanding platform and unraveled the potential roles of primary carbohydrate metabolism in sink tissues. As a result, we identified starch metabolism-related products, sugar and sugar alcohols, and TCA cycle intermediates as key molecular drivers of sink strength in poplar and summarized the major findings as a postulated model in Fig 7. The genes and metabolites identified in sink tissues in this study are potential targets for genetic engineering to improve sink strength in poplar plants. As patterns of ^13^C allocation to belowground differ between growth periods (in particular in trees) (Zhou et al. 2022), a time-course study should be performed to understand the dynamics of C allocation and partitioning throughout the early and later stages of sink tissue growth and development. To further complement our understanding of the C allocation process in trees, future studies should also consider the complexity of tree perenniality and design experiments in the field under various environmental perturbations. Furthermore, belowground C allocation, including root exudation and C transfer to microbial communities in the rhizosphere, can constitute an important C sink function (Hartmann et al. 2020) which is a very important and exciting topical area to be investigated in future studies.

### Data and Materials Availability

The datasets generated and analyzed during the current study will be made available in the respective data repositories shortly after this submission.

## Acknowledgments

We thank Dr. Stephen DiFazio (West Virginia State University) for providing poplar woody stem cuttings. We also thank Nathan Johnson (Graphic Design Professional) for his assistance in refining figures 1 and 7 in this manuscript.

## Authors’ Contributions

**Vimal Kumar Balasubramanian**: designed and performed experiments, contributed to data analyses, wrote the original draft, reviewed, and edited the manuscript; **Albert Rivas-Ubach**: performed metabolomics (GC- and LC-MS) analyses and contributed to data analyses, reviewed, and edited the manuscript; **Tanya Winkler**: performed experiments including hormonal treatments, reviewed, and edited the manuscript; **James Moran**: conducted C isotope labeling experiment and contributed to the data analyses, reviewed, and edited the manuscript; **Hugh Mitchell**: contributed to RNA seq. data analyses, reviewed, and edited the manuscript; **Amir H. Ahkami**: conceptualized the work, acquired funding, designed experiments, wrote and edited the manuscript.

## Supplementary data

**Figure S1.**
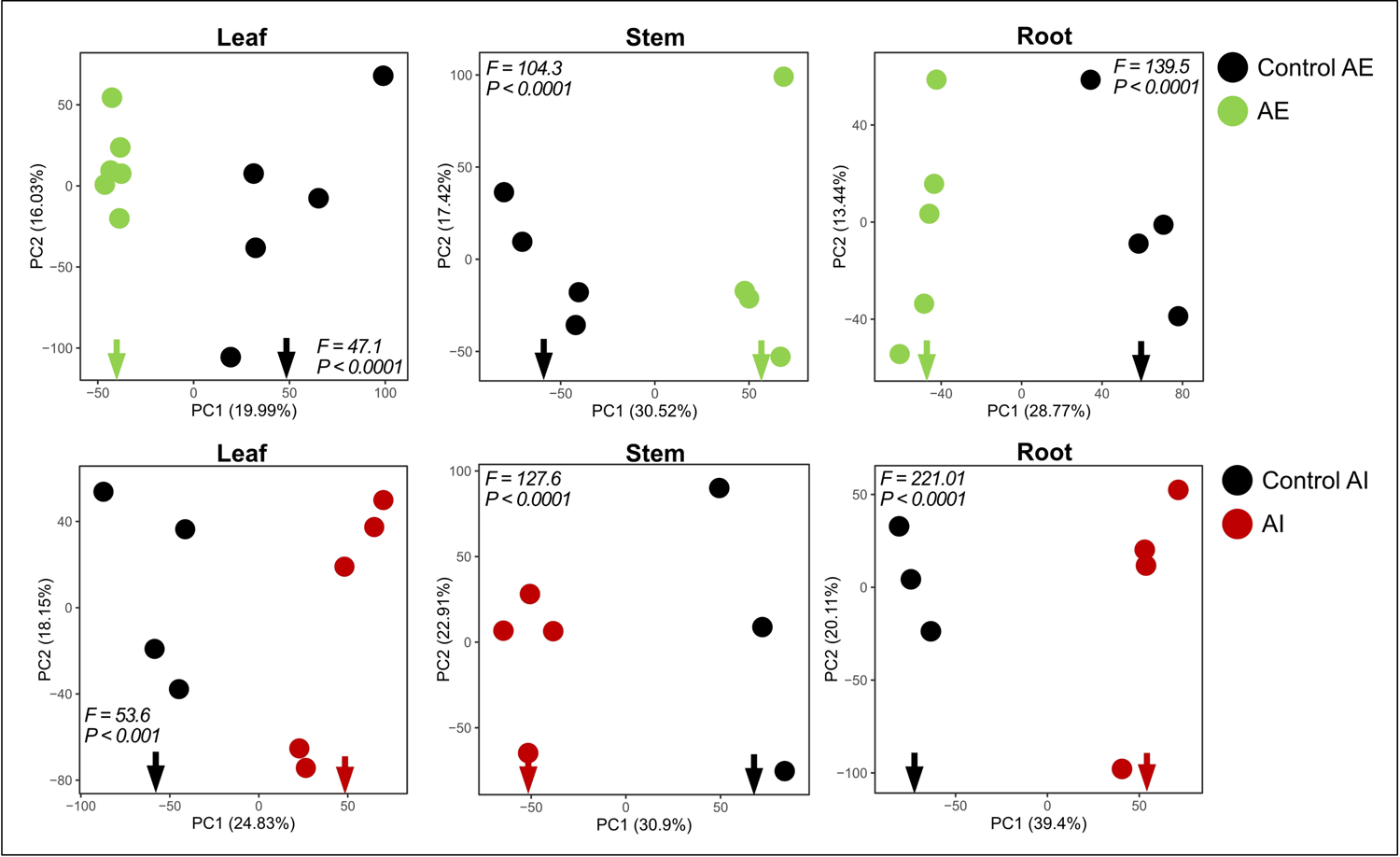
Principal Component Analysis (PCA) of total metabolomic profiles from leaf, stem, and root tissues in AE and AI foliar sprayed samples along with their respective control plant tissues.

**Figure S2.**
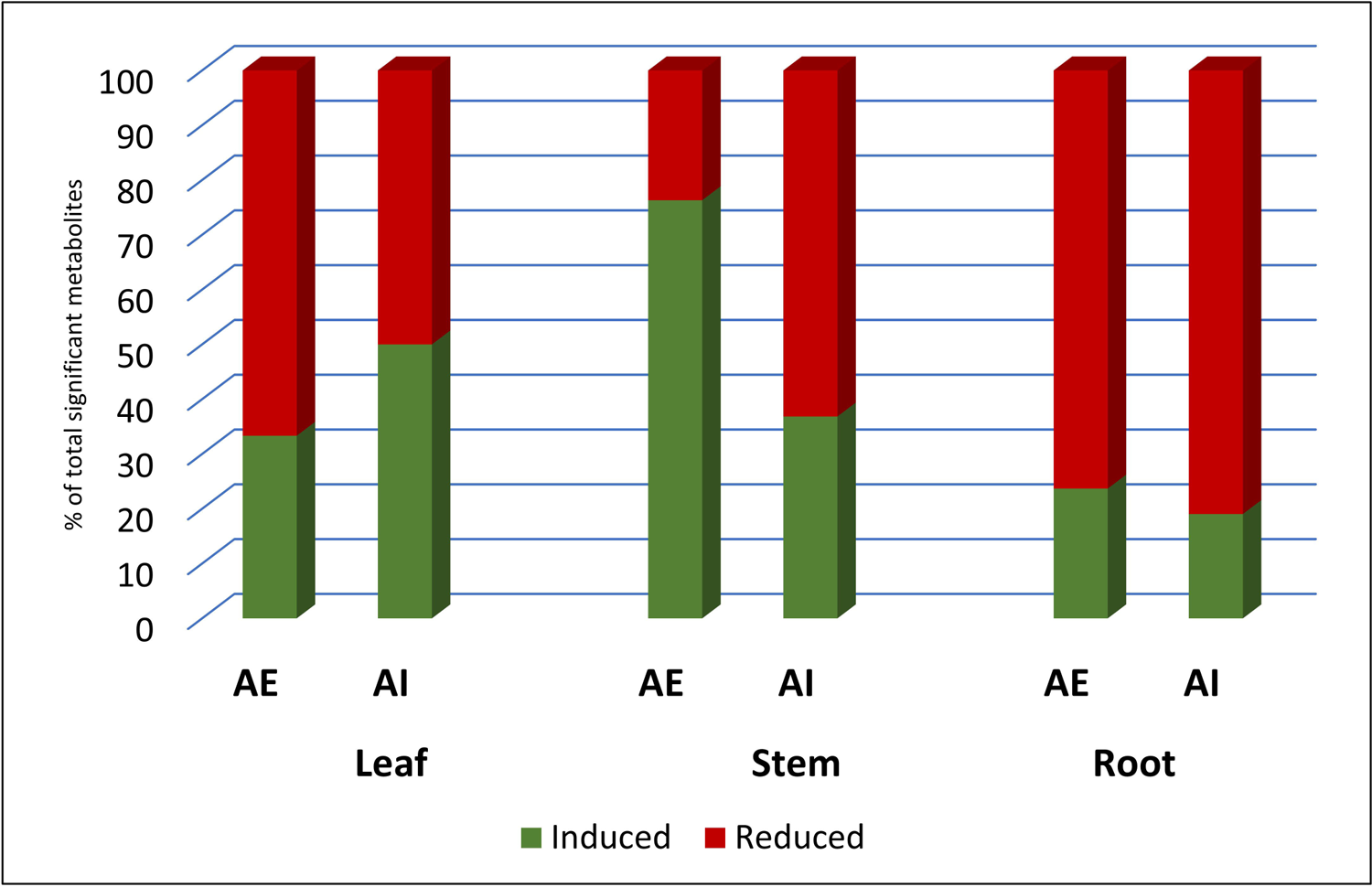
The total number of significantly altered metabolites in AE and AI-treated leaf, stem, and root tissues vs. their respective control tissues.

**Figure S3-A.**
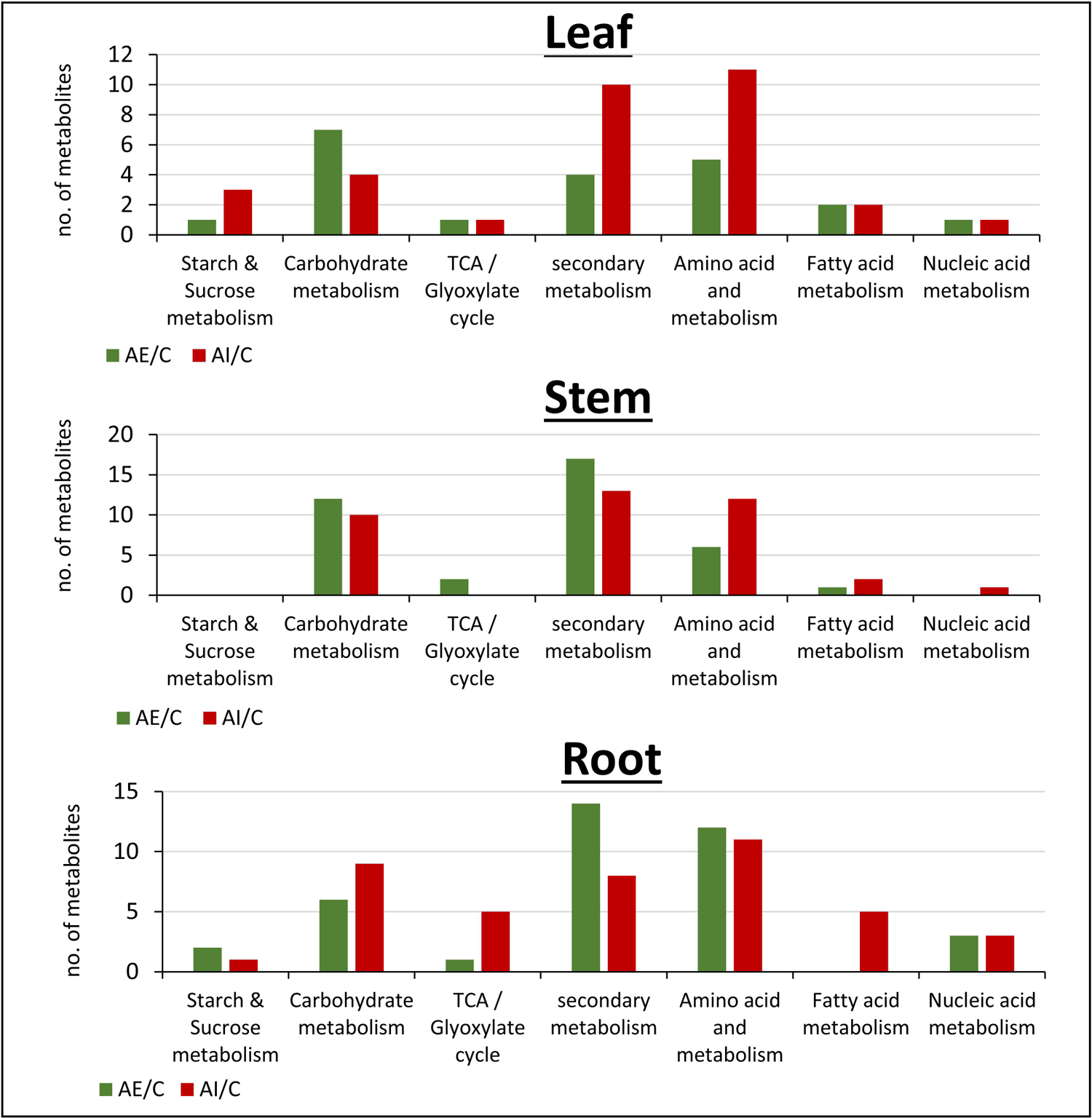
The total number of significant metabolites in AE and AI-treated leaf, stem, and root tissues vs. control tissues categorized into seven major KEGG pathway categories.

**Figure S3-B.**
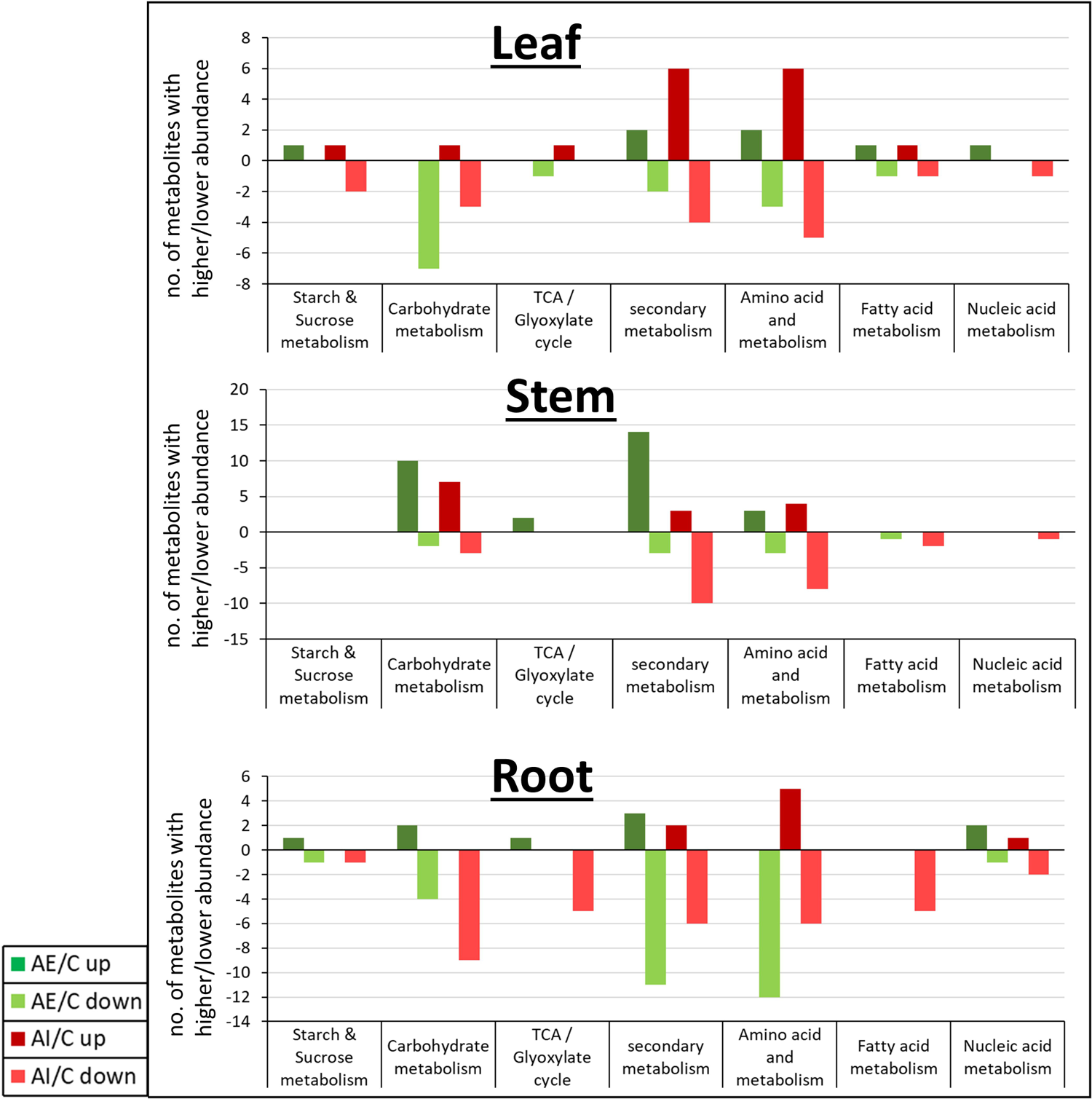
The total number of significant metabolites with higher and lower abundance levels in AE and AI-treated leaf, stem, and root tissues vs. control categorized into seven major KEGG pathway categories.

**Figure-S4-A.**
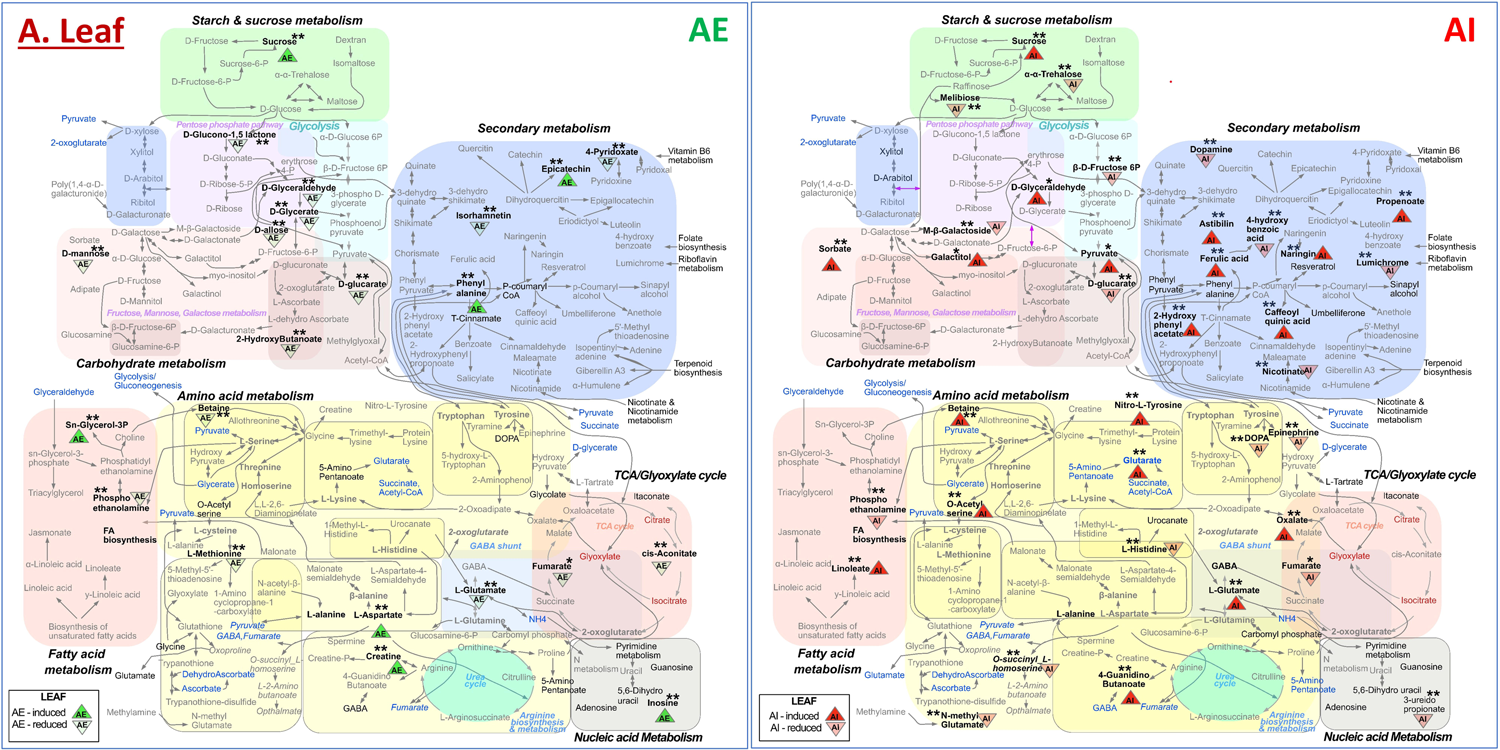
Metabolic pathway outline of leaf tissue under AE/AI foliar-sprayed and control conditions.

**Figure-S4-B.**
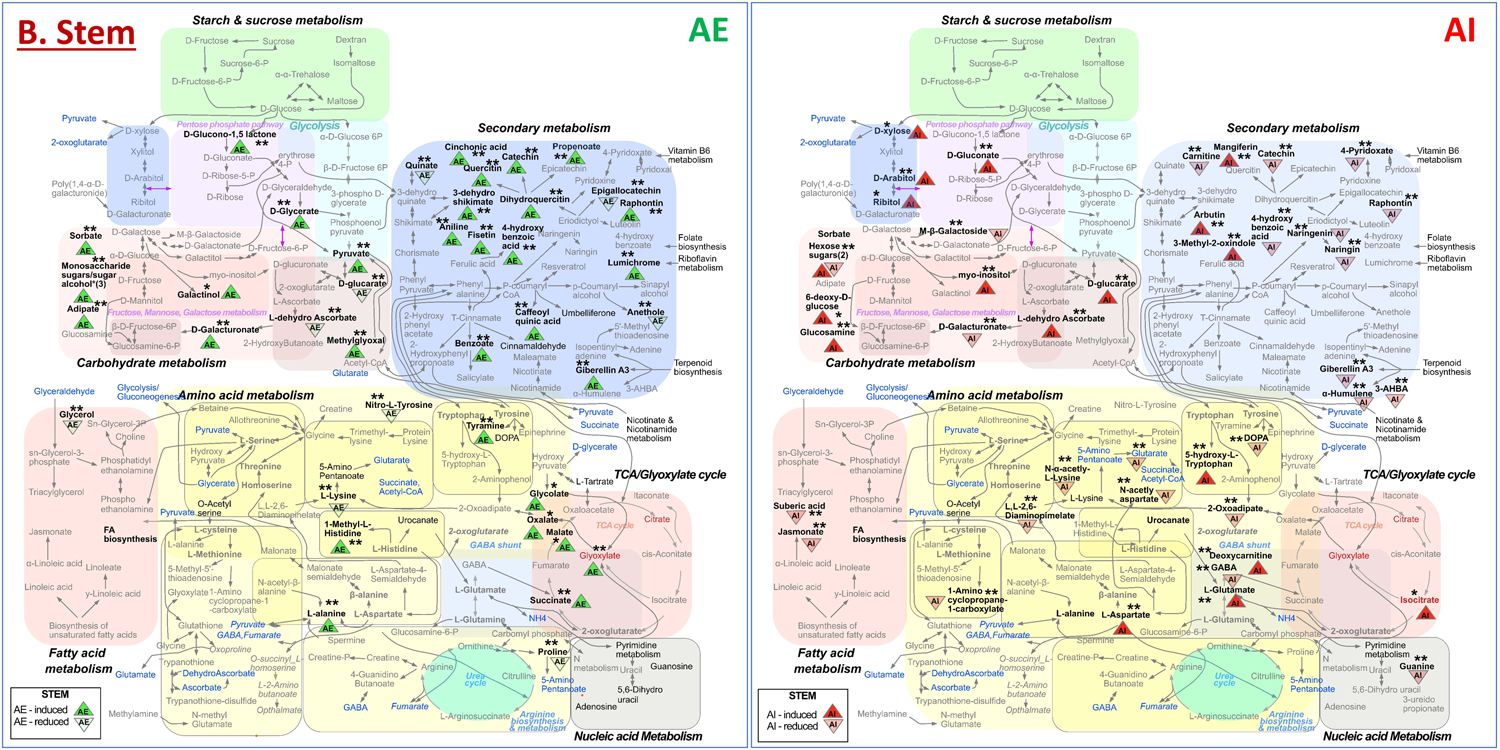
Metabolic pathway outline of stem tissue under AE/AI foliar-sprayed and control conditions.

**Figure-S4-C.**
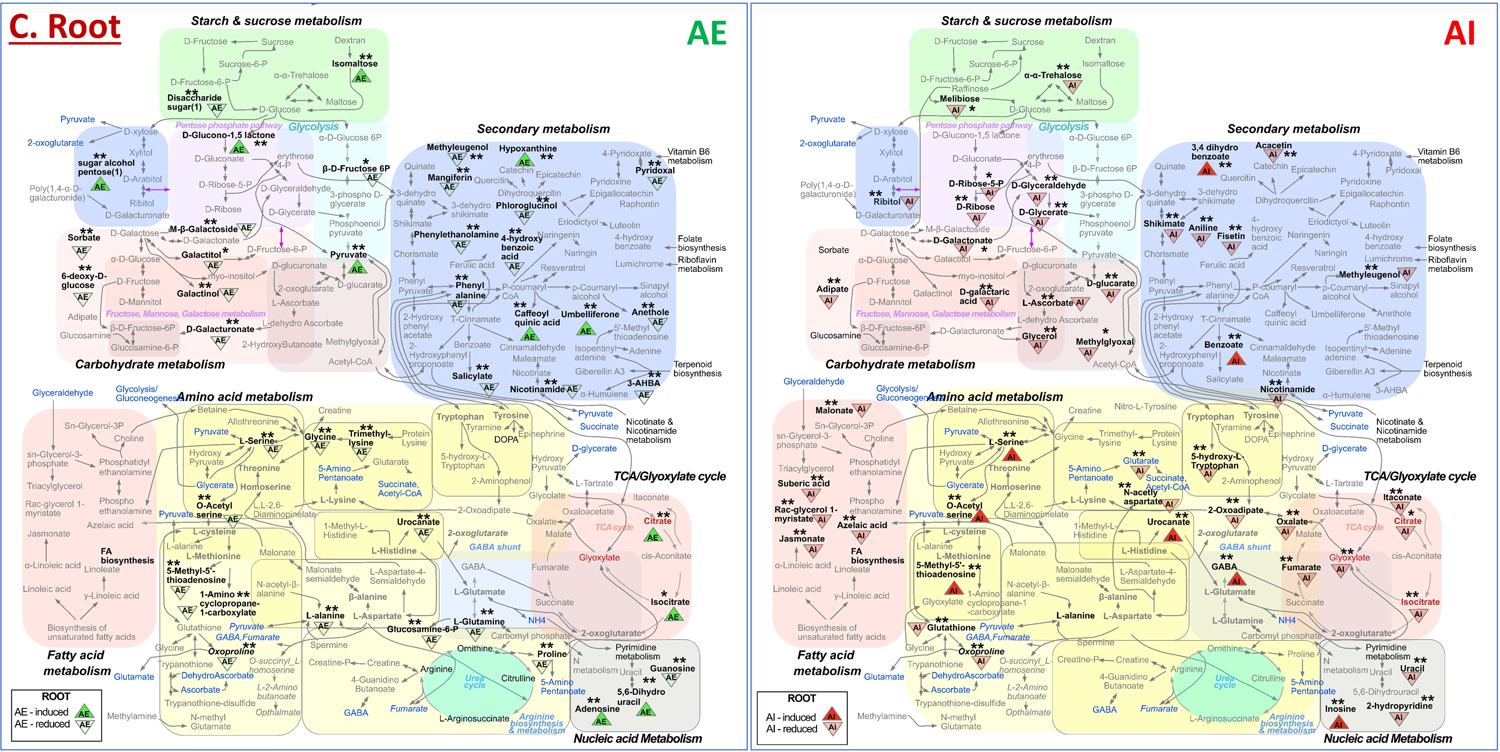
Metabolic pathway outline of root tissue under AE/AI foliar-sprayed and control conditions.

**Table S1:** Dry biomass weight data collected from control and hormone-(AE and AI) sprayed plant tissues (leaf, stem, and root) collected 11 days post spraying of hormones.

**Table S2:** Root image analysis and the values of selective derived parameters.

**Table S3:** Differential expressed genes from AE-treated plant roots vs. controls.

**Table S4-A:** PERMANOVA analysis highlighting significant differences on the overall metabolome of hormone-treated plant tissues vs. control plant tissues.

**Table S4-B:** Metabolite abundances from LC-MS and GC-MS analysis of auxin enhancer (AE) treated plant tissues (leaf, stem, and root) vs. control.

**Table S4-C:** Metabolite abundances from LC-MS and GC-MS analysis of auxin inhibitor (AI) treated plant tissues (leaf, stem, and root) vs. control.

**Table S4-D:** Parameters applied to GC-MS chromatograms with Metabolite Detector 2.5 for obtaining the GC-MS metabolomic profiling of AE, AI and control treated poplar plants.

**Table S4-E:** Parameters applied to LC-MS RAW files with MZmine 2.51 for obtaining the LC-MS metabolomic profiling of AE, AI and control treated poplar plants

**Table S5:** ^13^C isotope enrichment analysis performed using an isotope ratio mass spectrometer (IRMS) and the output data containing measured δ^13^C (‰) values.

